# A Bayesian Informative Shrinkage Approach for Large-scale Multiple Hypothesis Testing (BISHOT): with Applications in Differential Analysis of Omics Data

**DOI:** 10.1101/2025.09.11.675690

**Authors:** Ya Su, Mary Eunice Joy Z. Clark, Chi Wang

**Affiliations:** Department of Statistical Sciences and Operational Research, Virginia Commonwealth University, Richmond, VA 23284-3083, U.S.A.; Division of Cancer Biostatistics, Department of Internal Medicine and Markey Cancer Center, University of Kentucky, Lexington, 40536, U.S.A.

**Keywords:** Bayesian large scale hypothesis tests, non-central global local shrinkage, differential gene expression, sign-adjusted classification risk

## Abstract

A major goal of many omics studies is to identify differential features, e.g. differentially expressed genes, between experimental groups. When performing differential analysis for a given dataset, relevant information from another platform or species is often available. Incorporating such prior information can help identify features that show consistent differential patterns across platforms or species, which are more likely to reflect shared biological processes, and thereby enhance the robustness and generalizability of the findings. However, existing differential analysis methods typically analyze only the data from the current study and do not leverage prior knowledge about the magnitude or direction of changes from other platforms or species. We address this challenge, and the associated multiple testing problem, using a Bayesian framework that enables the incorporation of prior knowledge obtained from different platforms or species. We propose a new test statistic, Bayesian Credible Ratio (BCR), based on a heteroscedastic global local shrinkage prior, and a new multiple testing criterion, sign-adjusted FDR (SFDR), that emphasize information regarding the direction of the differentially features. We prove that BCR achieves the largest count of sign-based true positives among all legitimate SFDR-controlling methods. Simulation results offer numerical evidence of its advantage compared to an empirical Bayesian method. The approach is demonstrated through the analysis of RNAseq and single-cell RNAseq datasets.

## 1 Introduction

The statistical differential analysis plays an important role in many omics studies that involve two or more experimental groups. Various statistical methods have been developed to identify differentially expressed genes from RNAseq or single cell RNAseq (scRNAseq) (Finak et al., 2015; Love et al., 2014; Robinson et al., 2010; Smyth, 2005; Wu et al., 2013), differentially bound sites from ChIPseq (Ross-Innes et al., 2012), differentially accessible peaks from ATACseq (Gontarz et al., 2020), or differentially abundant proteins/metabolites from mass spectrometry (Huang et al., 2020; Li et al., 2019) between experimental groups.

When conducting the differential analysis for a given omics dataset, there is often relevant information available from another platform or species. Such prior information enables investigators to distinguish features that exhibit consistent differential patterns across platforms or species from those specific to a single platform or species. Because features that are consistent across platforms or species likely reflect shared underlying biological processes (Vitorino, 2024), prioritizing them can enhance the robustness and generalizability of the findings (Altenbuchinger et al., 2017; Wang et al., 2022). A few such examples are as follows:

- Example 1: Hohmann et al. (2016) performed an RNAseq experiment on a human acute myeloid leukemia cell line, MV4, to investigate changes in the gene expression profile with the treatment of BI-7273, a BRD9 inhibitor. In addition to this experiment, the investigators also performed an RNAseq experiment on a mice cell line, RN2, using the same treatment. Hohmann et al. (2016) found that *MYC* expression was downregulated in BRD4 inhibitor–treated cells in both human (MV4) and mouse (RN2) cell lines. This consistent suppression of *MYC* across species provides strong evidence that BRD4 plays a critical role in sustaining *MYC* transcription.
- Example 2: Angelidis et al. (2019) conducted a scRNAseq experiment that profiled the cell types and gene expressions in old and young mice to study lung aging. An analysis of interest is to identify differentially expressed genes in type-2 pneumocytes, one of the most prevalent cell types in the samples, between old and young mice. In parallel, the authors also conducted a bulk RNAseq experiment on type-2 pneumocytes, selected by flow cytometry sorting, from old and young mice. Angelidis et al. (2019) found significant agreement between the scRNAseq results and bulk RNAseq results, supporting the robustness of the age-related genes identified by scRNAseq.
- Example 3: Li et al. (2022) performed an ATACseq experiment to identify chromatin regions related to resistance to second-generation androgen receptor inhibitors, e.g. enzalutamide, by comparing enzalutamide-sensitive and resistant cell lines. Prior to this experiment, the investigators had already performed an RNAseq experiment on the same cell lines (Li et al., 2020). Li et al. (2022) focused on consensus genes that were both differentially expressed in RNAseq data and had differentially accessible promoter peaks in ATACseq data, thereby narrowing down the list of candidate genes and ultimately identifying a potential therapeutic target, GSTM2, for overcoming resistance to enzalutamide.

A common approach for leveraging prior information is to consider a two-step procedure, where the first step is to perform differential analysis on the current dataset alone to identify significant features, and the second step is to only keep the subset of significant features that are consistent with prior information from other platform/species (Chen et al., 2024; Li et al., 2022). However, since all features are analyzed together and treated equally in the first step, the significance levels of true consistently differential features can be reduced due to the presence of platform/species-specific differential features, e.g. a relatively small (but significant) effect claimed by both prior and current data, therefore consistent, can be declared as insignificant due to low signal noise ratio in the current data, e.g., in the analysis of MV4 cell line data in Section 5, the current standard approach fails to identify *MYC* which is claimed to be significant by BISHOT. In addition, downstream analyses, such as gene set enrichment analysis, may become complicated since ranking genes based on significance levels is difficult with the filtering based on prior information.

Despite the demand for a rigorous procedure to incorporate these prior information, most existing differential analysis methods perform the analysis based on the given dataset alone without allowing historical prior information to be supplied (Robinson et al., 2010; Smyth, 2005; Sun and McLain, 2012). Although leveraging historical data has been explored (Li et al., 2017, 2016), the focus has primarily been on stabilizing the estimation of mean and variance parameters, either by treating historical data as informative priors (Li et al., 2016) or by grouping genes with similar variations from historical data (Li et al., 2017). To our knowledge, there is a lack of methods that explicitly incorporate prior knowledge about the magnitude or direction of changes, e.g. fold changes in gene expression, between experimental groups.

Another potential area of research is regarding multiple testing error control, given the number of genes in expression data grows substantially. The property of false discovery rate (FDR) has been extensively studied in the literature (Benjamini and Yekutieli, 2001; Genovese and Wasserman, 2004; Storey, 2003, among others). Various methods consider alternatives to the traditional *p*-value based FDR control approach (Benjamini and Hochberg, 1995), by using some ‘local’ measure for evidence towards *H*_*ag*_ (Efron, 2008; Stephens, 2017; Sun and McLain, 2012), which quantifies individualized ‘significance’ under data heteroscedasticity and arguably enhances the power in large scale testing problems. More evidence regarding benefits of empirical Bayes based procedures in handling multiple comparisons can be found in Efron (2012), Kendziorski et al. (2003), Muralidharan (2010) and references therein. As a comparison method, we briefly describe Sun and McLain (2012) shortly.

For simplicity purpose, we will derive our method based on a linear regression model for RNAseq data. An extension to a more complicated model for scRNAseq data is provided in Section 6. For gene *g* (*g* = 1, …, *G*) in subject *i* (*i* = 1, …, *n*), let *Y*_*ig*_ be the normalized expression level in the form of log_2_(CPM + 1), where CPM denotes counts per million. We consider a classical regression model that allows for heteroscedasticity for *Y*_*ig*_:

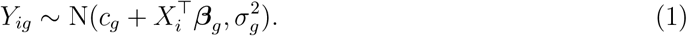

Generally speaking, *X*_*i*_ ∈ **R**^*p*^ is a set of covariates for subject *i, c*_*g*_ is the intercept, 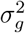 is the variance and ***β***_*g*_ is a vector of coefficients for gene *g*. The methodology is illustrated for the application of comparing RNAseq gene expressions between two groups, e.g. control and treatment, corresponding to model (1) with *p* = 1 where *X*_*i*_ ∈ *{*0, 1*}* represents a treatment group indicator, equivalently, *β*_*g*_ is the log_2_ fold change (LFC) quantifying the treatment effect for gene *g*.

The problem we are interested in is whether *β*_*g*_ is different from 0 for *g* = 1, …, *G*. A natural statistical formulation is via hypothesis test. Specifically we consider the null hypotheses to be composite, that is

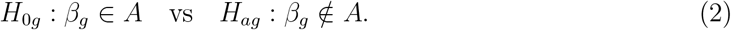

In this context we consider *A* = (−*ϵ, ϵ*), a small neighborhood containing 0. The composite null hypothesis ensures that identified differences are sufficiently large to be biologically meaningful, with the choice of *ϵ* informed by domain experts (McCarthy and Smyth, 2009).

A popular multiple hypothesis testing procedure is built upon some summary statistic which is normally distributed and centered on *β*_*g*_, that is, 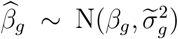. A common shrinkage prior with a zero component is typically utilized for *β*_*g*_, *g* = 1, …, *G*. That is, *β*_*g*_ ∼ *f* (*·*), with *f* (*x*) = *π*_0_*δ*_0_(*x*) + (1 − *π*_0_)*g*(*x*), where *δ*_0_(*x*) is a point mass at 0, *g*(*x*) is the distribution for non-zero signals and the probability of observing 0 is given by *π*_0_. Sun and McLain (2012) is specifically designed for composite null and the proposed ‘local’ estimate is proven to be optimal within a family of estimators called MLRC in large scale hypothesis testing.

There are several potential limitations in the above and similar procedures. First, they do not apply when there is prior knowledge about *β*_*g*_ as those procedures make heavy use of a common prior distribution for *β*_*g*_. Second, they rely on the existence of the summary statistics 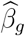 and its marginal distribution over *β*_*g*_, which are crucial in estimating *π*_0_ and *g*(*·*). Besides, estimations of *π*_0_ and *g*(*·*) are shown to be unstable from our simulation studies especially for smaller sample sizes.

We introduce a new **B**ayesian **i**nformative **s**hrinkage multiple **h**yp**o**thesis **t**esting (BISHOT) procedure that addresses the question of elucidating consistent features between two data sources with proven error control. Specifically, we propose a new shrinkage prior distribution where the local parameter plays a key role in governing the prior influence and is numerically shown to be adaptive to the feature-level coherence from prior to present. In addition, a new test statistic, Bayesian Credible Ratio (BCR), based on the posterior distribution is proposed which accounts for the heteroscedasticity of genes in favor of one side of tail over the other. While conducting multiple tests, we also take the tail-favoring perspective and propose a new criterion called the sign-adjusted FDR (SFDR). The calculation of BCRs for all genes combined naturally promotes parallelization across all genes. The decision process according to BCR is proven to attain the maximum number of true positives among all valid sign-adjusted false positive controlled procedures.

To the best of our knowledge, it is a difficult task for the available approaches to work with a more complicated model or when a vector ***β***_*g*_ is present due to the lack of explicit distribution of 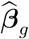 as well as multivariate deconvolution methods in the presence of unknown point mass on zero. The fully Bayesian framework we propose lays out a potential path to this setting without the need of establishing 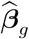 and carefully modeling *f* (*·*) and thus it is free from estimating non-null probability and non-null distribution. We present our solution for a two component model for scRNAseq data in Section 6.

Outline of this paper is as follows. Section 2 describes the construction of the prior distribution and the test statistic named the Bayesian credible ratio, its connection with the decision rule. Section 3 introduces a classification risk-based framework for large-scale differential analysis, detailing threshold selection and establishing the procedure’s optimality. Section 2 and Section 3 combined lead to the proposed BISHOT. Section 4 includes simulation results and comparisons with the above popular method. Applications to RNAseq and scRNAseq gene expression data are illustrated in Section 5 and 6. We conclude with several discussions in Section 7.

## 2 Bayesian credible ratio

We illustrate the proposed method using model (1), we later dive into a more complicated model applicable to scRNAseq data in Section 6. The likelihood function corresponding to (1) for a given gene feature *g* along with observed expressions *i* = 1, …, *n* is:

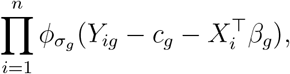

by convention, *ϕ*_*σ*_ denotes the normal density with mean zero and standard deviation *σ*.

We propose a heteroscedastic global local shrinkage (HGLS) prior for *β*_*g*_ based on our knowledge about *β*_*g*_,

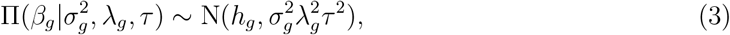

the input of non-central parameter *h*_*g*_ is reflective of the information given about *β*_*g*_, in this work, *h*_*g*_ is a fixed value and set as the prior estimate for the treatment effect associated with feature *g* based on findings from earlier studies. The prior variance consists of the local parameter *λ*_*g*_ and the global parameter *τ*, in addition to the standard deviation *σ*_*g*_ from error distribution in (1).

HGLS for *β*_*g*_ builds upon the classical global and local prior by explicitly modeling the substantial heteroscedasticity present in gene expression data. Contrary to zero, it has a shrinkage effect towards *h*_*g*_, the existing knowledge about *β*_*g*_. The shrinkage is also relative to the variability of that specific gene feature captured by *σ*_*g*_. The local shrinkage *λ*_*g*_ governs the prior influence and is learned in a data-driven manner based on the gene-specific coherence between historical and current data. As illustrated in Figure 5, *λ*_*g*_ increases with the discrepancy between historical and current treatment effects, thereby reducing the impact of historical values when they are not aligned with current findings.

In this work we adopt standard prior distributions for shrinkage parameters *λ*_*g*_ and *τ* from horseshoe. That is, *λ*_*g*_, *τ* ∼ Half-Cauchy(0, 1). The heteroscedastic error variance 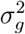 takes a conjugate prior, 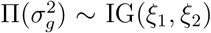 with *ξ*_1_ and *ξ*_2_ set according to the moments of empirical estimates of 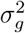 given the observed gene expressions. The overall expression *c*_*g*_ is assigned a prior distribution 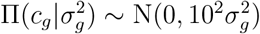 in the general case.

Under the proposed prior (3) and model (1) for the expression data, we can derive the (conditional) posterior distribution of 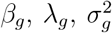 and *τ* given data *D*_*g*_ = *{Y*_*ig*_, *i* = 1, …, *n}* and *h*_*g*_, where *λ*_*g*_ and *τ* are sampled using slice sampling algorithm, similar to Bhattacharya et al. (2016). For better mixing of the chain, we sample *β*_*g*_ given *λ*_*g*_ and *τ* while integrating out *σ*_*g*_. The conditional posterior distributions for all parameters are explicitly written in the Appendix A.

The proposed test procedure is based on the *tail evidence* from the posterior distribution of the targeting parameter. For the purpose of illustration, in the remainder of the section we simplify the notation from *β*_*g*_ and *D*_*g*_ to *β* and *D* correspondingly, e.g., posterior distribution of *β* is denoted as Pr(*β*|*D*).

- We define the Bayesian credible ratio (BCR) for the hypothesis test *H*_0_ : *β* ∈ (−*ϵ, ϵ*) vs *H*_*a*_ : *β* ∈*/* (−*ϵ, ϵ*) by *C*_*β*_ = *C*_0,*β*_(1 − *C*_*a,β*_)^−1^, where C_0,*β*_ = Pr(*β* ∈ (−*ϵ, ϵ*)|*D*) and C_*a,β*_ = min*{*Pr(*β* ≤ −*ϵ*|*D*), Pr(*β* ≥ *ϵ*|*D*)*}* are the posterior probability for *β* over (−*ϵ, ϵ*) and its smaller tail over (−∞, −*ϵ*) or (*ϵ*, ∞).
- The null hypothesis is rejected if *C*_*β*_ is smaller than a threshold 0 ≤ *λ* ≤ 1.

Given the model and the prior structures, the hypothesis test in (2) is analyzed by BCR based on the posterior samples of *β*_*g*_. BCR compares evidence towards the null/alternative hypotheses by *C*_0,*β*_ and 1 −*C*_*a,β*_. The notion of BCR is model and prior free and might be of interest for its own purpose. When hypotheses regarding all feature effects are considered, we suggest the threshold *λ* be selected to control the sign-adjusted FDR (SFDR) in the multiple testing regime, see details in Section 3.

### 2.1 BCR as a Bayesian decision rule

Generally speaking, a hypotheses testing problem *H*_0_ : *β* ∈ Θ_0_ vs 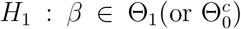 can be formulated as a Bayesian decision process (Berger, 2013). Let *a*_0_/*a*_1_ be the action of accepting of *H*_0_/*H*_1_, the loss function for *a*_*i*_ is *L*(*a*_*i*_, *β*), *i* = 0, 1. Consider a simple ‘0 − *K*_*i*_’ loss function, *L*(*a*_*i*_, *β*) = 0 if *β* ∈ Θ_*i*_ and *L*(*a*_*i*_, *β*) = *K*_*i*_ if *β* ∈*/* Θ_*i*_. The expected posterior loss for *a*_0_ and *a*_1_ is *K*_0_*P* (Θ_1_|*D*) and *K*_1_*P* (Θ_0_|*D*) correspondingly. The Bayes rule opts for the action which minimizes the expected posterior loss, that is, accepting *H*_0_ when *P* (Θ_0_|*D*) *> K*_0_*/*(*K*_0_ + *K*_1_) and accepting *H*_1_ otherwise.

When Θ_0_ = (−*ϵ, ϵ*), we revise the above decision process as follows. First since the alternative set 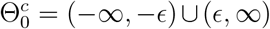 is composed of two non-overlapping sets, we consider performing two hypotheses with alternatives being *H*_1_ : *β* ∈ Θ_1_ = (−∞, −*ϵ*) and *H*_2_ : *β* ∈ Θ_2_ = (*ϵ*, ∞). In this circumstance, we define three actions *a*_0_, *a*_1_ or *a*_2_ corresponding to accepting of *H*_0_, *H*_1_ or *H*_2_. Suppose the ‘0 − *K*_*i*_’ loss functions are chosen for *a*_*i*_, *i* = 0, 1, 2 under a constraint *K*_1_ = *K*_2_ (no distinction of loss under *a*_1_ or *a*_2_), the Bayes rule regarding *H*_0_ and *H*_1_ is accepting *H*_0_ when *P* (Θ_0_|*D*)*K*_1_ *> K*_0_*P* (Θ_1_|*D*) and accepting *H*_1_ otherwise; the Bayes rule regarding *H*_0_ and *H*_2_ is accepting *H*_0_ when *P* (Θ_0_|*D*)*K*_1_ *> K*_0_*P* (Θ_2_|*D*) and accepting *H*_2_ otherwise.

We propose the ‘universal’ Bayes rule regarding *H*_0_ : *β* ∈ Θ_0_ vs *H*_*a*_ : *β* ∈ Θ_1_ ∪ Θ_2_ as accepting *H*_0_ when both decisions made above agree with action *a*_0_ and accepting *H*_*a*_ when either of the above decisions says otherwise. Equivalently, the universal decision process is accepting *H*_0_ when *P* (Θ_0_|*D*) *>* (*K*_0_*/K*_1_) max*{P* (Θ_1_|*D*), *P* (Θ_2_|*D*)*}* and accepting *H*_1_ otherwise. Define *λ* = *K*_0_*/*(*K*_0_ + *K*_1_), 0 *< λ <* 1, our decision can be further formulated as accepting *H*_0_ if *C*_*β*_ *> λ* or accepting *H*_1_ if *C*_*β*_ *< λ*, where *C*_*β*_ is the BCR defined previously.

## 3 Optimal decision rule for differential analysis via sign-adjusted classification risk

In the presence of a large number of features, the issues of multiple testing emerge when hypotheses tests (2) are carried out for each feature individually. To design an optimal decision rule while controlling the marginal false discovery rate (mFDR), Sun and Cai (2007) shows the equivalence to study the weighted classification risk corresponding to *β* ∈ (−*ϵ, ϵ*) or *β* ∈*/* (−*ϵ, ϵ*). The optimal decision rule is shown to be the minimizer to the classification risk, while the associated statistic is optimal in that it minimizes the false nondiscovery rate, subject to a constraint on the false discovery rate.

As detailed in Section 2.1, the Bayes decision rule governed by BCR, is based on two separate decisions regarding *β*_*g*_ ≤ −*ϵ* or *β*_*g*_ ≥ *ϵ*. We will show that this Bayes rule is an estimator of the optimal decision/minimizer to a weighted classification risk with sign preferences for all features. Furthermore, it is proven to capture the maximum number of sign-adjusted expected true positives while controlling for the marginal SFDR (mSFDR). The proofs to all Theorems in this Section can be found in the Appendix A.

### 3.1 BCR as minimizer of classification risk utilizing sign preferences

Define

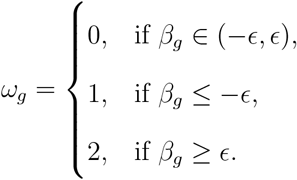

We purposely treat *β*_*g*_ ≤ −*ϵ* and *β*_*g*_ ≥ *ϵ* separately since the combination of the data and the informative shrinkage prior should provide evidence (if any) favoring one of them rather than both.

Define *δ*_*g*_ = 1 if the null hypothesis for feature *g* is rejected and 0 otherwise, ***δ*** and ***ω*** represent the concatenated vector of *δ*_*g*_ and *ω*_*g*_ for *g* = 1, …, *G*. Let *λ* ∈ [0, 1] represent the weight governing the false positive/negative losses. Define 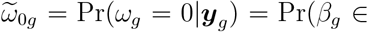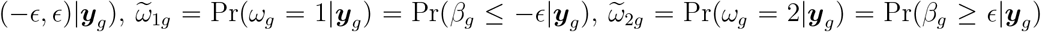, where ***y***_*g*_ = *{Y*_1*g*_, …, *Y*_*ng*_*}* is the vector of observations for feature *g*.

Consider first two loss functions to losses of detecting negative effects (*ω*_*g*_ = 1) or positive effects (*ω*_*g*_ = 2) correspondingly:

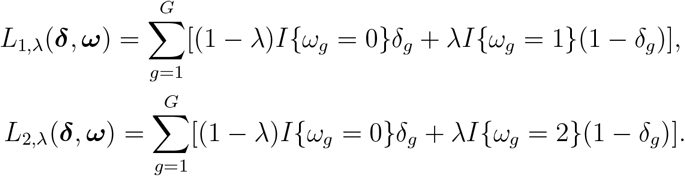

Theorem 1. For *k* = 1, 2, the minimizer of *EL*_*k,λ*_ (the expected value of *L*_*k,λ*_) is denoted by ***δ***_*k*_, which is the vector form of *{δ*_*kg*_, *g* = 1, …, *G}* where 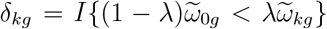. *EL*_*k,λ*_ is defined over the probability law of ***δ*** and ***ω***.

Next, we define a partition of all features with regards to the preference of positive or negative signs when rejecting. This partition exists under Assumption 1 below. Incorporating information about the sign of the parameter to construct decision, or to choose between the terms in *L*_1,*λ*_ or *L*_2,*λ*_, leads to a more conservative decision than ignoring the sign. It is preferred in our setting when features are provided with additional confidence per prior knowledge (3).

**Assumptions 1**. For a fixed 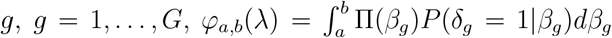, either (i) or (ii) holds: (i) *φ*_*ϵ*,∞_(*λ*) ≤ *φ*_−∞,−*ϵ*_(*λ*) for all *λ* ∈ (0, 1) (ii) *φ*_*ϵ*,∞_(*λ*) ≥ *φ*_−∞,−*ϵ*_(*λ*) for all *λ* ∈ (0, 1).

Assumption 1 requires the sign preference is invariant to *λ*, that is, case (i) and case (ii) holds universally for all any choice of *λ. φ*_−∞,−*ϵ*_(*λ*) has an equivalent form 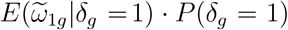 and similarly 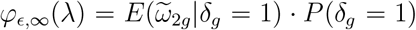. Therefore, under case (i) or (ii) 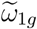 is expected to be larger or smaller when the decision is rejection.

Let 𝒮 or 𝒮^*c*^ denote the set of features which belong to case (i) or (ii) correspondingly. Based on Assumption 1, loss in *L*_1,*λ*_ tends to concern features in 𝒮 than those in 𝒮^*c*^, this motivates us to prioritize *ω*_*g*_ = 1 or *ω*_*g*_ = 2 in each scenario and thus combine ***δ***_1_ and ***δ***_2_ into one. The combined decision we propose is 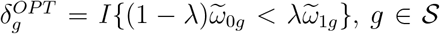 and 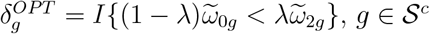. Denote ***δ***^*OPT*^ as the concatenated vector.

#### Theorem 2.

***δ***^*OP T*^ minimizes *EL*_3,*λ*_ where

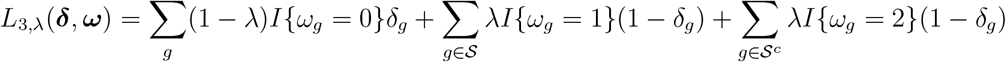

The optimal decision ***δ***^*OP T*^ has one missing piece: the set 𝒮 is unknown and needs to be estimated. When *δ*_*g*_ = 1, it is expected that 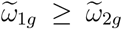 if *g* ∈ 𝒮 and 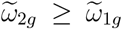 if *g* ∈ 𝒮^*c*^. Therefore the optimal decision can be estimated empirically as 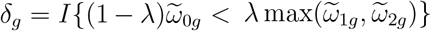 . According to the definitions of 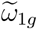 and 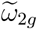, one comes to an equivalent expression 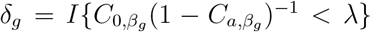 where 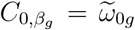 and 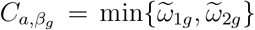. This establishes the relationship that BCR is an estimator of 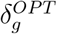.

### 3.2 Selection of *λ* in the multiple testing regime

The choice of *λ* can be selected by controlling the SFDR at the desired level *α*. In practice,

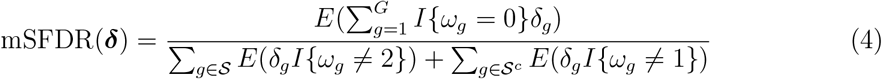

which denotes the marginal SFDR, and it can be shown that mSFDR = *SFDR* + *O*(*G*^−1*/*2^) using the same technique as Genovese and Wasserman (2002) for the asymptotic equivalence for mFDR and FDR. mSFDR is the mFDR based on sign of the parameters and has a natural link to the classification risk *L*_3,*λ*_.

Although the explicit form of mSFDR(***δ***) is generally unknown, we can estimate it using the following procedure (a slight modification of Section 3.2 in Sun and McLain (2012)):

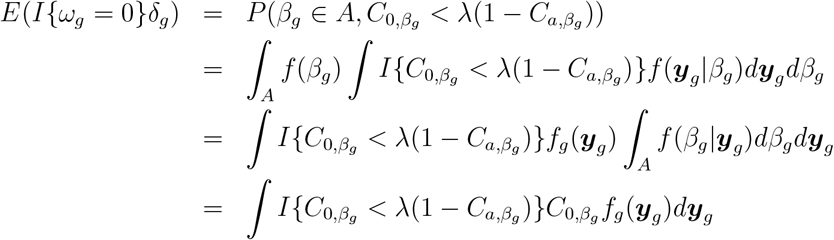

The numerator of mSFDR(***δ***) can be unbiasedly estimated by 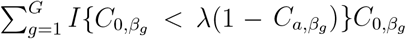. The same arguments hold true for the denominator of mSFDR(***δ***), which can be estimated by 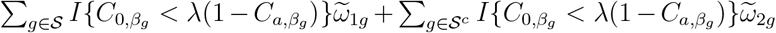. Together with these estimators, we select *λ* ∈ (0, 1) to be the largest such that 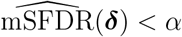.

### 3.3 Optimal property of *δ*^*OP T*^

The discovery of true positives is one criterion to compare procedures given the same FDR *α*. Under additional knowledge about the sign of the signal (given rejection), as the proposed mSFDR, the true positives count is modified accordingly, that is, true positives count are refined with information about sign appropriately.

#### Definition 1.

The sign-adjusted expected number of true positives is defined as follows:

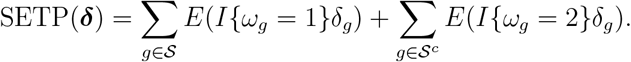

Clearly, compared to the traditional ETP, SETP only counts when the parameter *β*_*g*_ is in the correct sign given rejection, which is divided up by either 𝒮 or 𝒮^*c*^ (Remark 3.1). The following Theorem states that the proposed decision is optimal in achieving the greatest SETP under the same mSFDR threshold.

#### Theorem 3.

Assume that *λ* is chosen such that mSFDR = *α* for a fixed *α*. The procedure ***δ***^*OP T*^ has the largest SETP among all valid mSFDR procedures.

## 4 Simulations

### 4.1 Simulation setup

Our simulation studies is based on model (1) which can be useful for comparing gene expressions in two groups, e.g., control and treatment. This is achieved by setting *X*_*i*_ to be an indicator (0 or 1) variable if *i*th expression in the treatment group, *β*_*g*_ thus becomes the treatment effect.

We select a simulation scheme where 100(1 − *p*)% of the genes share the same expression mechanisms between two groups. That is, *β*_*g*_ is set to zero except for 100*p*% of the genes that are randomly selected. The proportion *p* is normally referred to as the proportion of differentially expressed (DE) genes. The following choices are used *p* = 0.1 or *p* = 0.5 corresponding to low or moderate presence of DE genes. Denote the set of DE genes as Ω, hence *β*_*g*_ = 0 when *g* ∈*/* Ω. For *g* ∈ Ω, *β*_*g*_ ∼ N(log_2_(2), 0.1^2^) (up to a random sign placement). In the Appendix A, a separate simulation with expressions of DE genes generated from a unimodal distribution is conducted.

In model (3) the following values are used: *c*_*g*_ = 4, *σ*_*g*_ = 1 for a total *G* = 5000 genes. The sample size for treatment/control group is *n* = 10 or *n* = 50 with the total sample size for the two groups doubled. The rest of the hyperparameters required are set as follows. The hyperparameters *ξ*_1_ and *ξ*_2_ are determined such that the first and second moments for all (pooled) sample variances of the expressions (combining treatment and control group) match with those under the prior. We consider two options for *ϵ* in the hypothesis tests (2), log_2_(1.5) or log_2_(1.2) targeting at moderate or small treatment effects in DE analysis. The prior distributions for *β*_*g*_ is centered on *h*_*g*_ if *g* ∈ Ω. For *h*_*g*_, we purposely perturb the true value of *β*_*g*_ with *h*_*g*_ ∼ N(*β*_*g*_, 0.5^2^), that is, the information regarding *β*_*g*_ is only within a vicinity of its true value. We generate *N* = 50 replicated data sets under each scenario and results are summarized in Section 4.2.

We provide a concise description of the method that is compared to BISHOT, denoted as SunSpike0. As mentioned in the introduction, the algorithm in Sun and McLain (2012) is theoretically-justified and suitable for composite null hypothesis. Since it relies on estimating the distribution of *β*_*g*_ with a zero component and a distribution for DE genes given 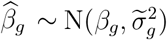, it is not directly applicable to any data-generating model (when the distribution of 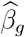 does not have an explicit form as such) but can be adapted to model (1) if 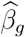 is selected to be the MLE and 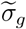 as the standard error. To the best of our knowledge the code underlying Sun and McLain (2012) is not publicly available. Alternatively, the `R` package `ashr` (Stephens, 2017) can extract the distribution of *β*_*g*_ assuming the distribution for DE genes is unimodal and symmetric with which we estimate the test statistic in Sun and McLain (2012). The resulting estimate then used in controlling mSFDR is compared with BISHOT in the simulation above. Similar findings are present in simulation II where the unimodal and symmetric distribution is used as the true distribution for *β*_*g*_ (the Appendix A).

### 4.2 Simulation results

Under a collection of SFDR nominal values ranging over [0, 0.25], we validate the actual SFDR achieved using (4) with *δ*_*g*_ and the true *ω*_*g*_. Figure 1 and 2 display the actual SFDR values averaged over 50 simulations against the corresponding nominal SFDR thresholds for the four scenarios given by *n* and *p* when *ϵ* = log_2_(1.2) (Figure 1) and *ϵ* = log_2_(1.5) (Figure 2). It can be seen that almost always BISHOT produces SFDR much closer to its nominal value while SunSpike0 tends to yield much higher FDR and not surprisingly the advantage becomes more prominent when the sample size is smaller and/or there is less DE genes (small *n* and/or small *p*). It is noteworthy that SunSpike0 behaves quite differently when the proportion of DE genes *p* and sample size *n* changes, conversely, BISHOT is dramatically more stable/robust among all scenarios given, thanks to the informative prior. Generally speaking, *ϵ* has an impact on the conservativeness of both methods, genes with small or zero *β*_*g*_ become more distinct from the genes with large *β*_*g*_ relative to a larger threshold hence the actual false discovery proportion tends to decrease when *ϵ* increases. Hence the desired conservativeness of the method can provide some guidance of choosing *ϵ*.

**Figure 1:**
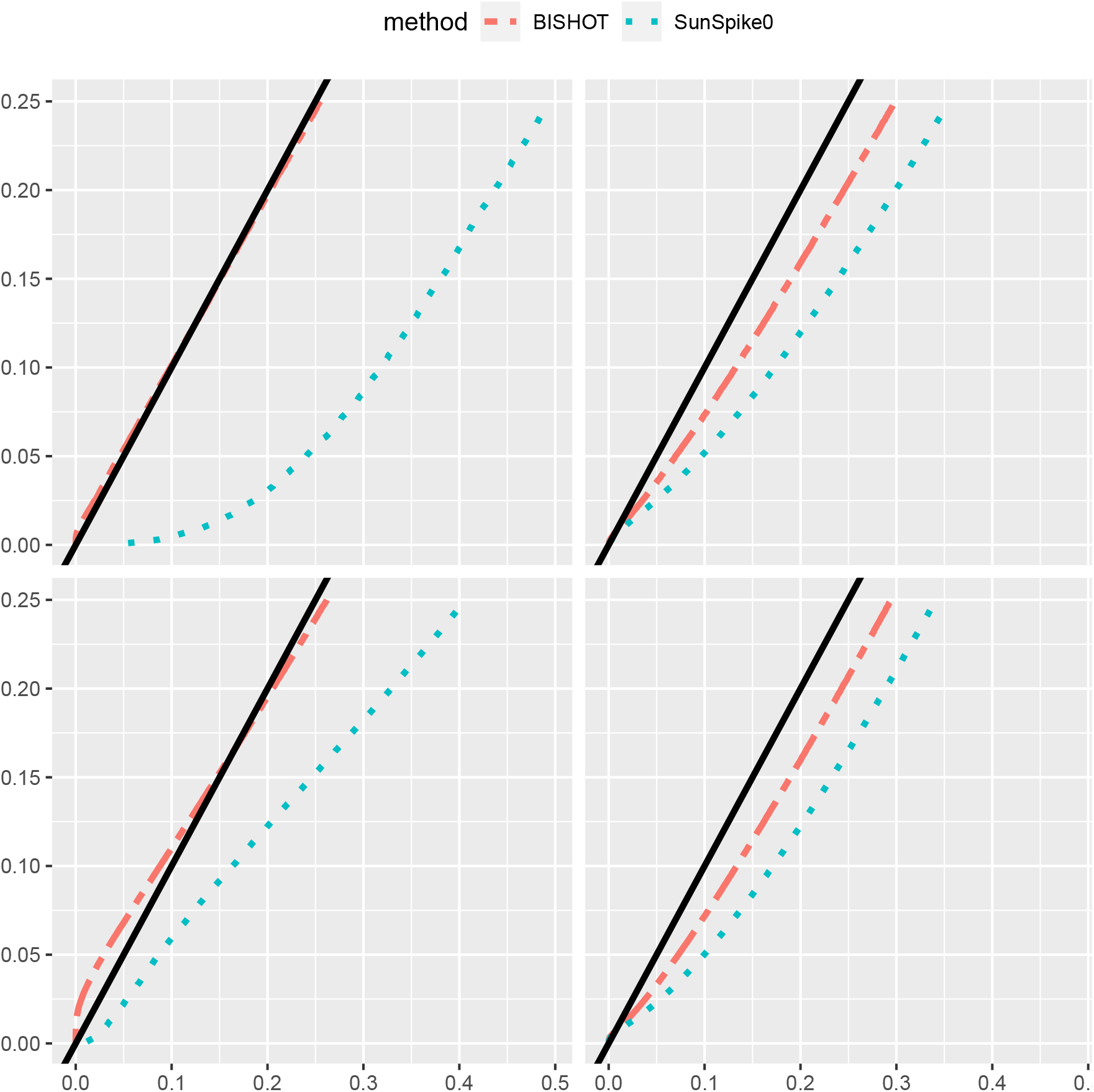
The actual signed false discovery rate (SFDR) when *ϵ* = log_2_(1.2) averaged over 50 simulated data using mSFDR threshold 0.001 to 0.25 corresponding to 10 (left panel) or 50 (right panel) samples in the treatment and control group. The top and bottom panel displays the case when *p* = 0.1 and *p* = 0.5. The solid black line corresponds to the *y* = *x* line.

**Figure 2:**
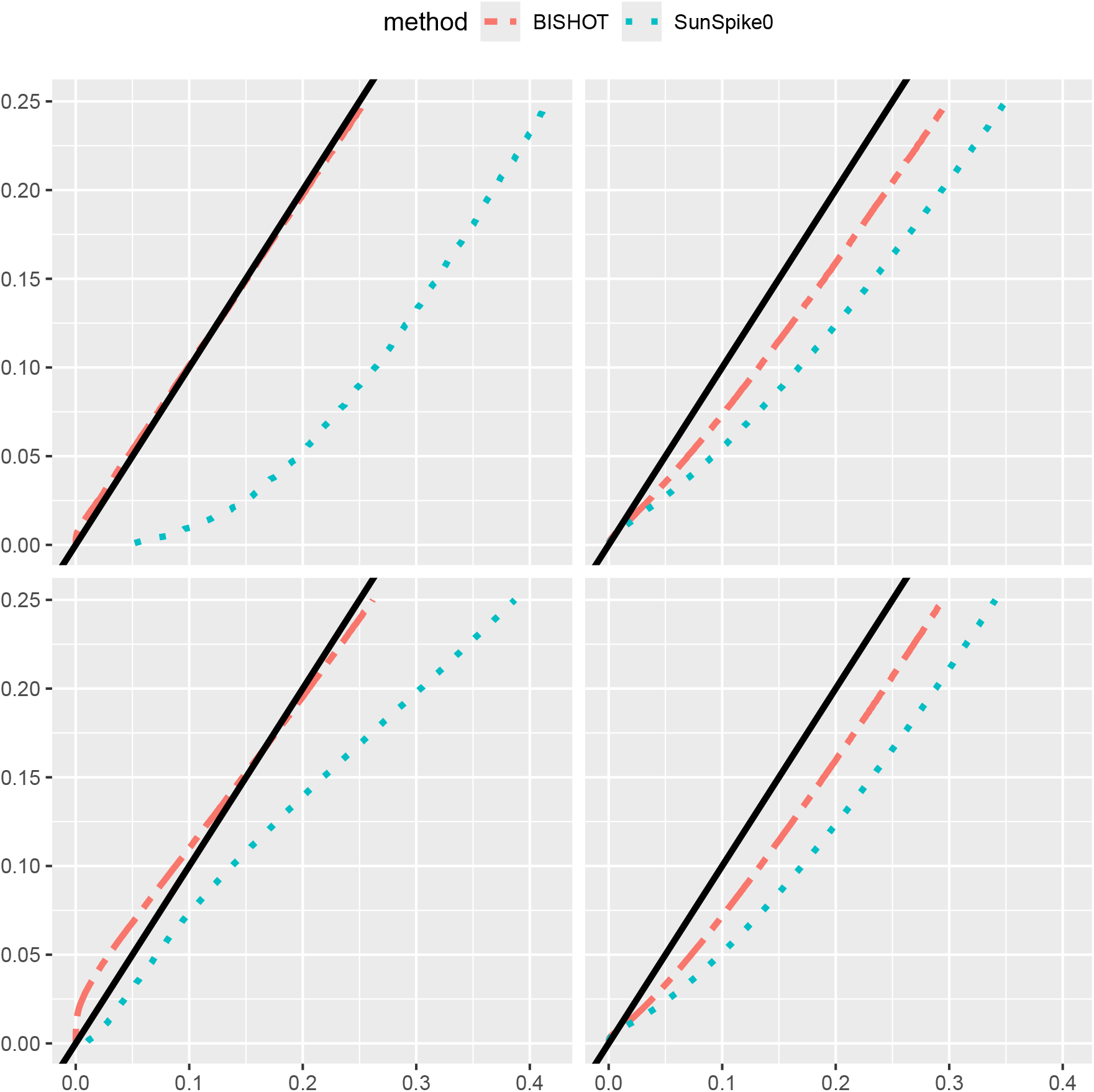
The actual signed false discovery rate (SFDR) when *ϵ* = log_2_(1.5) averaged over 50 simulated data using mSFDR threshold 0.001 to 0.25 corresponding to 10 (left panel) or 50 (right panel) samples in the treatment and control group. The top and bottom panel displays the case when *p* = 0.1 and *p* = 0.5. The dashed black line corresponds to the *y* = *x* line.

We next compare how the top-ranked genes match the true DE genes. For this purpose, we select the first *R* genes, 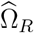, ranked by their SFDR based *q* values (the minimum SFDR you can achieve by calling a given test significant) and report the proportion of those fall in Ω whose absolute values of *β*_*g*_ are greater than *ϵ* as well, that is, 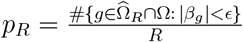. The value *R* is selected from 1 to the total number of DE genes plus 100, that is, *Gp* + 100. The averaged proportion *p*_*R*_ for all simulations against *R* is shown in Figure 3 and 4 corresponding to *ϵ* = log_2_(1.2) and *ϵ* = log_2_(1.5). The *R* highest ranked genes by BISHOT are never falsely detected for even relatively large *R* (relative to the size of Ω), while SunSpike0 does not perform as good especially when expression sample size is small regardless of *p* and *ϵ*. Hence BISHOT is more robust and their top ranked genes are more accurate.

**Figure 3:**
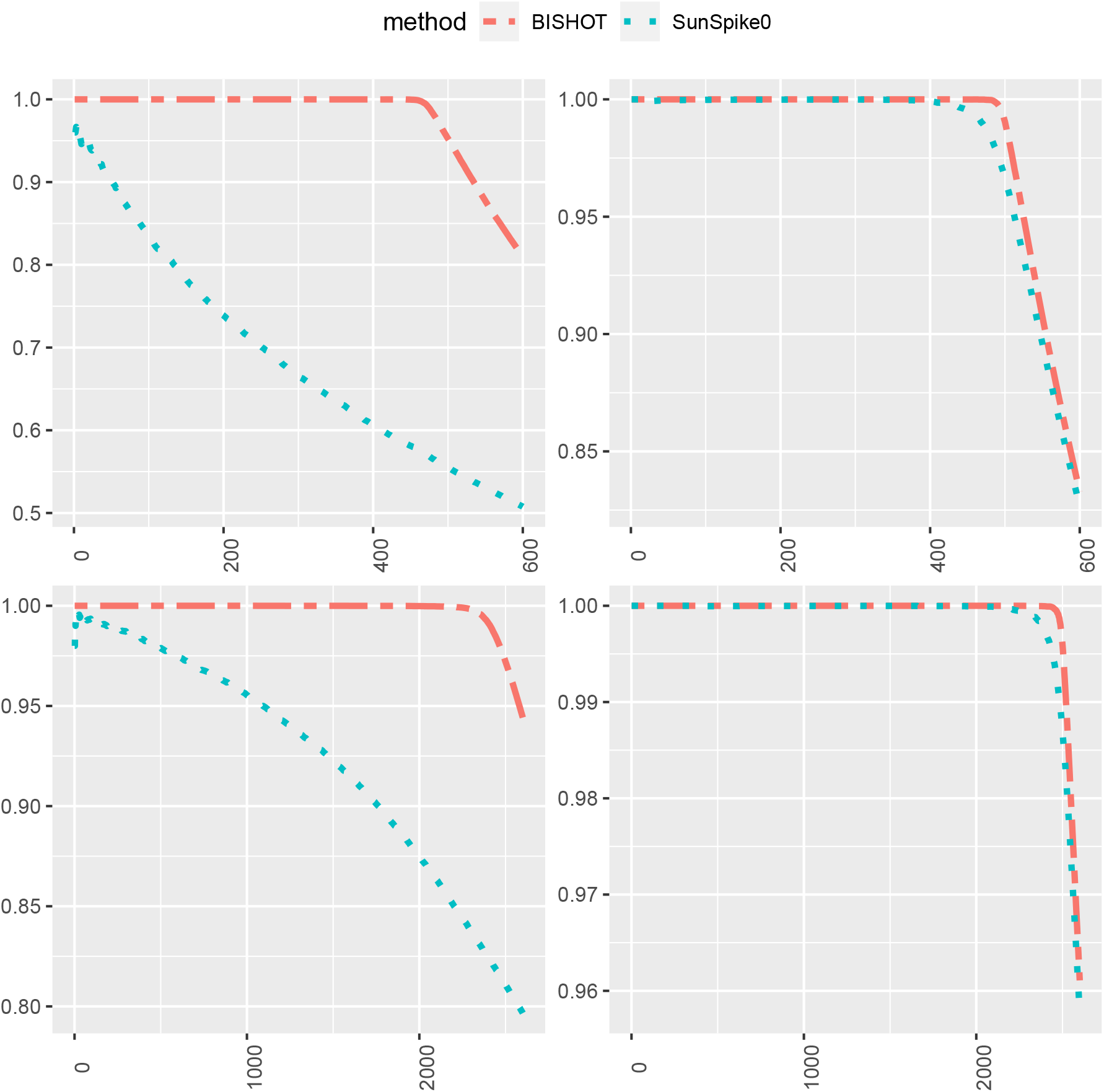
The proportion of true DE genes for the first *R* genes with smallest *q* values when *ϵ* = log_2_(1.2) averaged over 50 simulated data for *R* = 1, …, 5000*p* + 100 corresponding to 10 (left panel) or 50 (right panel) samples in the treatment and control group. The top and bottom panel displays the case when *p* = 0.1 and *p* = 0.5.

**Figure 4:**
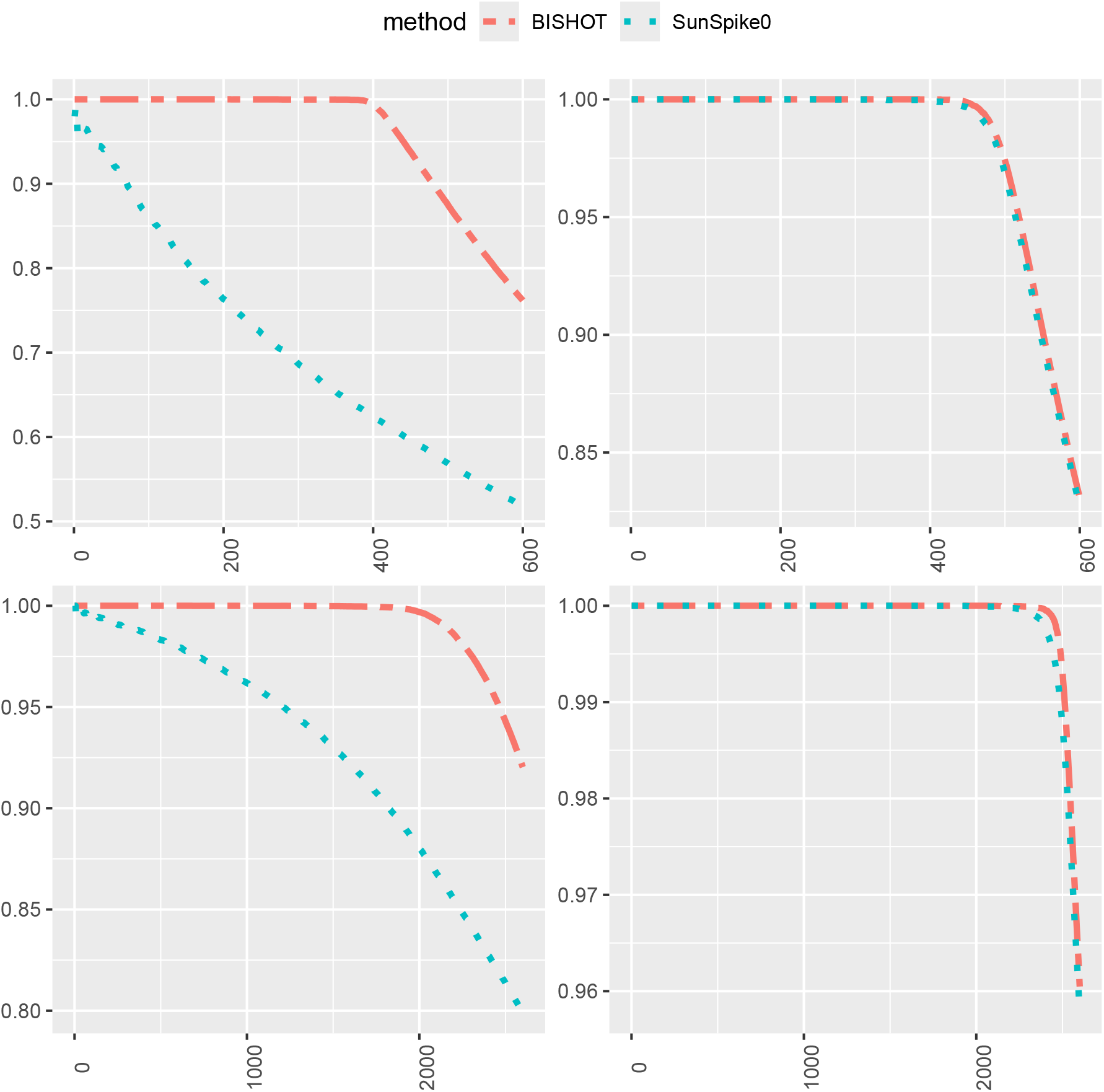
The proportion of true DE genes for the first *R* genes with smallest *q* values when *ϵ* = log_2_(1.5) averaged over 50 simulated data for *R* = 1, …, 5000*p* + 100 corresponding to 10 (left panel) or 50 (right panel) samples in the treatment and control group. The top and bottom panel displays the case when *p* = 0.1 and *p* = 0.5.

## 5 Transcriptional effects of BRD9 inhibition in acute myeloid leukemia

Acute myeloid leukemia (AML) is a type of blood cancer with a five-year survival rate of 31.9% (Surveillance Research Program, National Cancer Institute, 2025). Hohmann et al. (2016) investigated the potential of BRD9, a subunit of the SWI/SNF chromatin remodeling complex that promotes cell proliferation, to serve as a therapeutic target for AML. Hohmann et al. (2016) conducted two RNAseq experiments, one using murine RN2 cell line and the other using human MV4 cell line. In each experiment, cells were treated with BI-7273, a BRD9 inhibitor, or control with two replicates per group. RNAseq data were generated to elucidate the transcriptional effects of BRD9 inhibition.

We applied BISHOT to the Hohmann RNAseq data, which were transformed into log_2_(CPM + 1) values. We focused on identifying DE genes from human MV4 cell line while using the RNAseq data from murine RN2 cell line as prior knowledge. Specifically, the parameter *h*_*g*_ was specified based on the LFC for gene *g* in the data from the murine RN2 cell line. Two options for *ϵ* are considered, log_2_(1.2) or log_2_(1.5), same as what are used in Section 4.2.

In Figure 5 we illustrate the effect of shrinkage by examining the relationship of the local shrinkage *λ*_*g*_ with the consistency between prior knowledge based on RN2 and what the MV4 data conveys about *β*_*g*_, measured by the distance between *h*_*g*_ and the maximum likelihood estimator 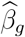. The shrinkage is in accordance to this consistency, to be more specific, it is prominent when 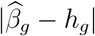 is small and less so when 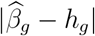 is large.

**Figure 5:**
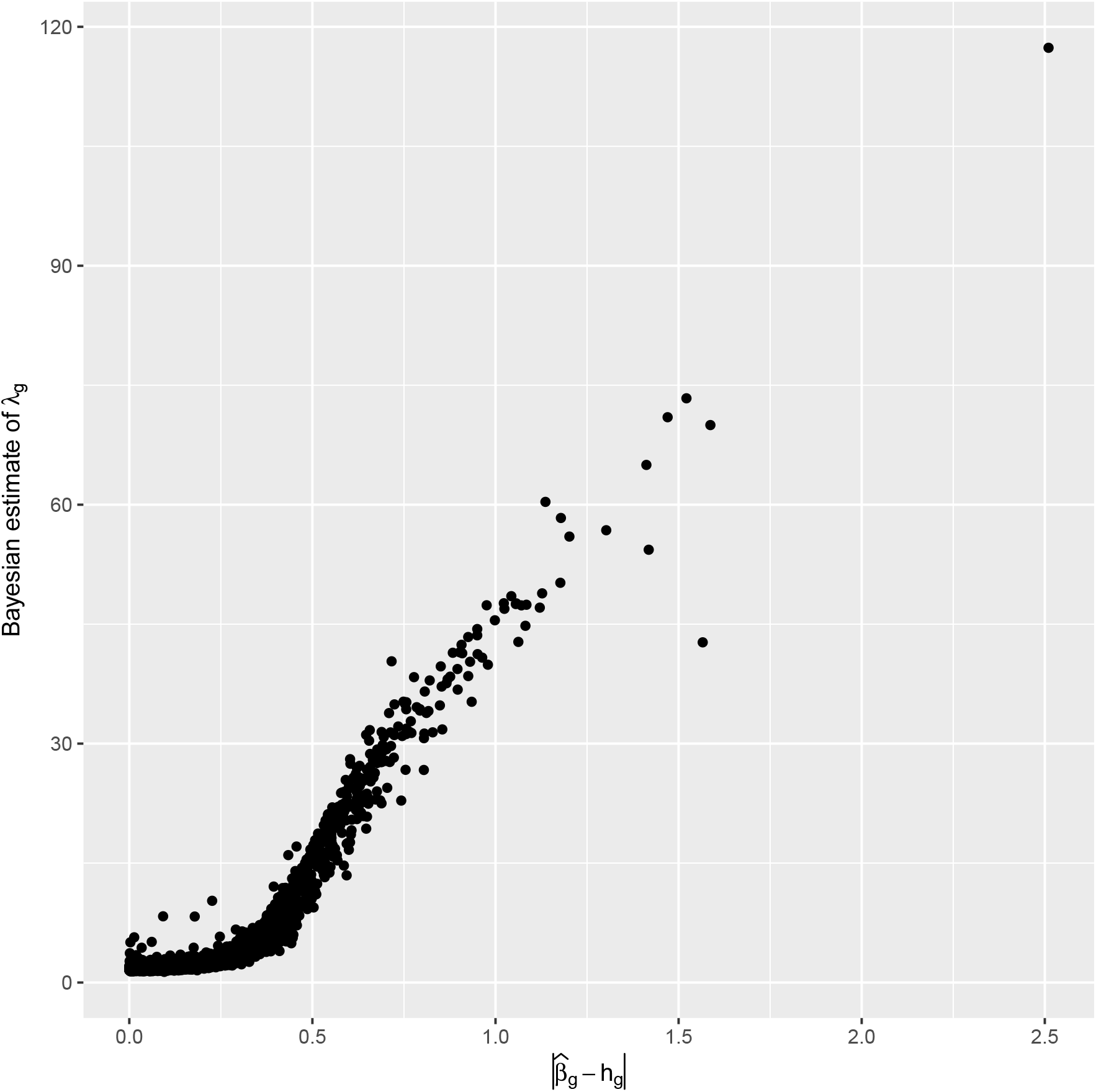
The relationship between the gene-specific shrinkage parameter and the distance between the two sources of information. The former is estimated by the posterior mean and the latter is by the absolute value of the difference between the Maximum likelihood estimator and the center of the prior distribution.

Figure 6 presents the identified DE genes at SFDR *<* 0.05 based on BISHOT relative to LFCs present in the RN2 and MV4 data. Almost all the genes with consistently large absolute LFCs (*> ϵ*) in both RN2 and MV4 (top-right and bottom-left regions) were identified as DE, except for a few genes with high variations. Conversely, no genes with consistently small LFCs (*< ϵ*) in both RN2 and MV4 (middle region) were identified as significant. Hence, by sharing information, the prior data from RN2 reinforced the analysis of MV4 data in determining DE genes, thanks to the reduction of posterior uncertainty when information is consistent.

**Figure 6:**
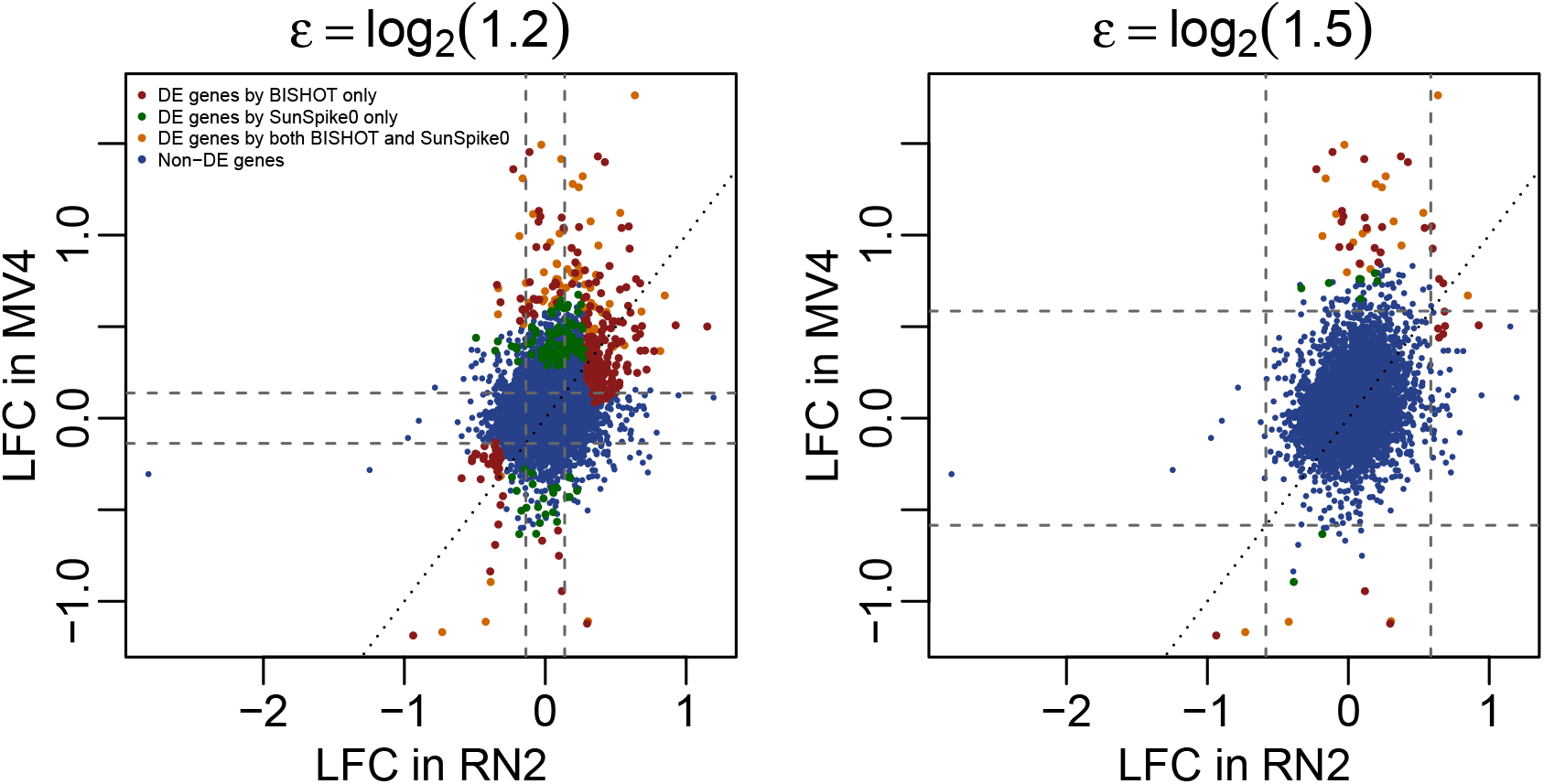
Gene expression LFCs between drug-treated and control groups based on RN2 (x-axis) and MV4 (y-axis) cell lines. Each dot represents a gene, colored according to DE results from the MV4 cell line. Left panel: *ϵ* = log_2_(1.2); right panel: *ϵ* = log_2_(1.5).

For genes with large LFCs in MV4 but relatively small LFCs in RN2 (vertical band outside the middle square), many were also identified as DE, demonstrating that our method prioritizes genes strongly supported by observed data, even when prior information is limited. This is important, as differences between prior data (e.g., mice) and the data under analysis (e.g., human) are expected, and the method should be able to detect features specific to the current dataset. Interestingly, many genes with large LFCs in RN2 but relatively small LFCs in MV4 (horizontal band outside the middle square) were not identified as differentially expressed, likely due to inconsistent LFCs or high variability in MV4. This result highlights that prior information alone is insufficient. Genes must be supported by the data under analysis to be considered significant.

Figure 6 also highlights the difference in DE results between BISHOT and SunSpike0. Genes that were called as DE by SunSpike0 only (green dots) tend to be the ones with absolute LFCs *> ϵ* in the MV4 but not in the RN2 data. In contrast, genes that were called as DE by BISHOT only (red dots) tend to be the ones with LFCs consistently greater than *ϵ* in both MV4 and RN2 data. In other words, BISHOT preferentially identifies signals that exhibit consistency between the two datasets. A Venn diagram comparing the number of DE genes between BISHOT and SunSpike0 is provided in the Appendix B.

We also compared the results from BISHOT with other popular methods available in `R` including limma (Smyth, 2005), voom (Law et al., 2014), and edgeR (Robinson et al., 2010). The limma (using the `lmFit` function in the `limma` package with log_2_(CPM + 1) values as the input), voom (using the `voomLmFit` function in the `limma` package with raw counts as the input), and edgeR (using the `edgeR` package with raw counts as the input) methods did not identify any DE genes at FDR *<* 0.05.

A key gene of interest in Hohmann et al. (2016) is *MYC*, where the authors showed that BRD4 plays a critical role in sustaining *MYC* transcription. We compared the significant level of *MYC* gene across different differential analysis methods. BISHOT with an *ϵ* = log_2_(1.2) yielded a *q*-value of 0.02, whereas all other methods produced non-significant *q*-values (Figure 7). Because *MYC* had a moderate LFC of −0.33 in the MV4 data, it was missed by differential analysis methods relying solely on that dataset. However, because *MYC* also showed a LFC of −0.36 in the RN2 data, which was highly consistent with that in the MV4 data, BISHOT with an *ϵ* = log_2_(1.2) was able to leverage such prior information to identify the consistent down-regulation of *MYC*. In addition, we noted that BISHOT with an *ϵ* = log_2_(1.5) produced a non-significant *q*-value for *MYC* because the method emphasizes genes with large fold changes and therefore did not capture *MYC* given its moderate fold change.

**Figure 7:**
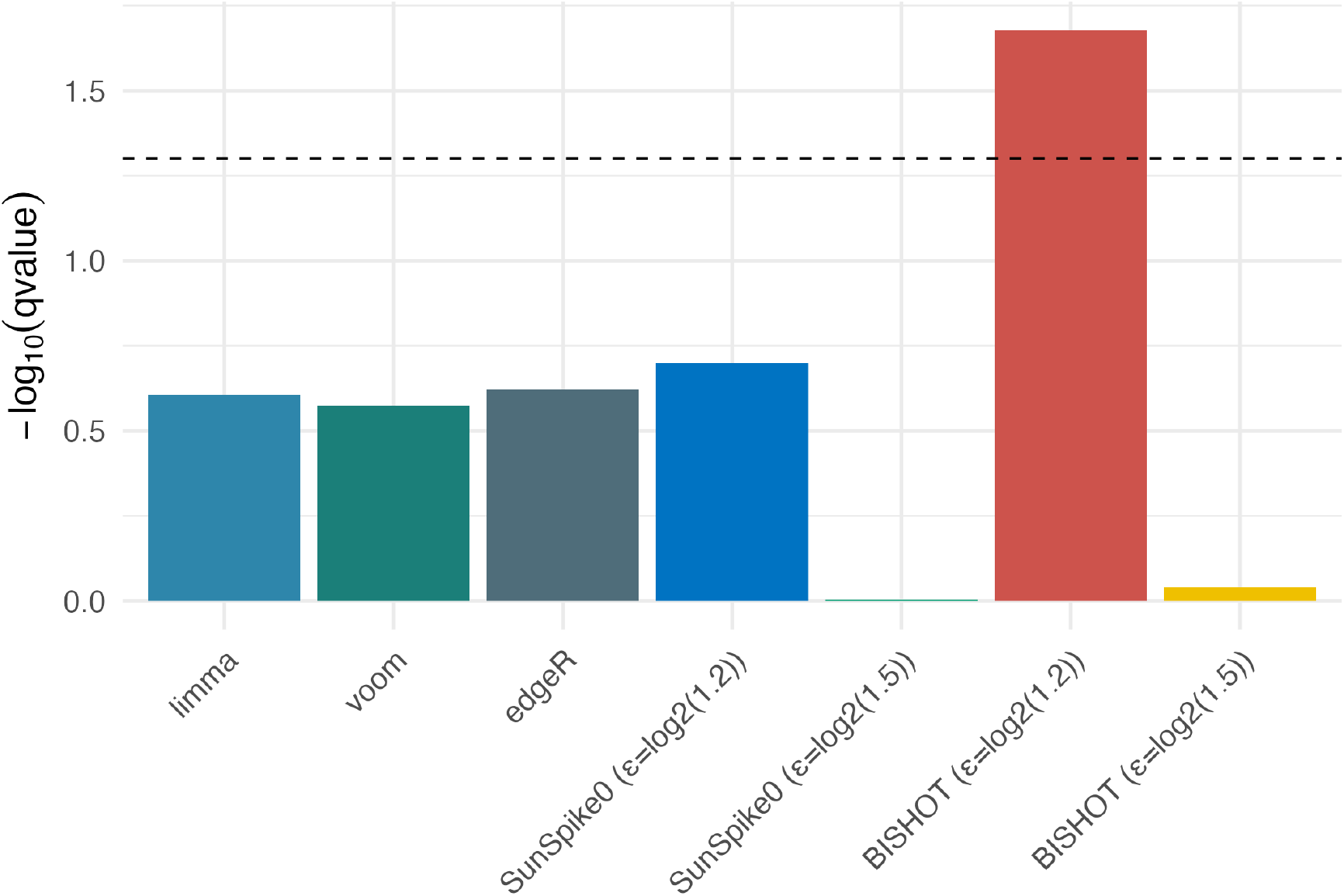
Comparison of the *q*-value for *MYC* across different methods. The dashed horizontal line indicates *q*-value=0.05.

## 6 The scRNAseq analysis of the aging lung

Angelidis et al. (2019) conducted a scRNAseq study in old (24 months) and young (3 months) mice to investigate cell type and gene expression alterations in lung aging. One of the most prevalent cell types identified in the study was type-2 pneumocytes. In parallel, the authors also conducted a bulk RNAseq experiment on type-2 pneumocytes, selected by flow cytometry sorting, from old and young mice. We applied BISHOT to scRNAseq data for type-2 pneumocytes to identify differentially expressed genes between old and young mice. The flow-sorted bulk RNAseq data for type-2 pneumocytes were used as our prior knowledge.

We first extend our model to handle scRNAseq data, which contains dropouts. Let *Y*_*ig*_ be the log_2_(CPM+ 1) expression value of gene *g* in cell *i*. Suppose *Z*_*ig*_ is a dropout indicator, where *Z*_*ig*_ = 1 if *Y*_*ig*_ ≠ 0 and *Z*_*ig*_ = 0 if *Y*_*ig*_ = 0. We adopt the two-component model developed in Finak et al. (2015): a logistic regression for *Z*_*ig*_ and a classic regression model for *Y*_*ig*_ conditioning on *Z*_*ig*_ = 1,

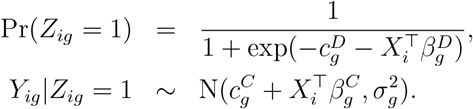

where corresponding to the probability component and the nonzero component, the parameters are superscripted by *D* or *C*. One would identify two treatment effects 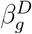 and 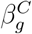 representing differentiations in terms of log odds for the probability of nonzero and the expected nonzero expression.

The question becomes which genes should be claimed significant based on either *β*^*D*^ or *β*^*C*^. Equivalently, we aim at a bivariate hypothesis for each gene, *H*_0*g*_ : ***β***_g_ ∈ ***A*** vs *H*_*ag*_ : ***β***_g_ ∈*/* ***A*** with 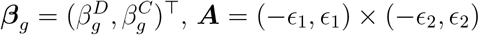. The choices of *ϵ*_1_ ∈ *{*log(1.2), log(1.5)*}* and *ϵ*_2_ ∈ *{*log_2_(1.2), log_2_(1.5)*}* correspond to a LFC or log odds ratio of 1.2 or 1.5 converted to the scale of *Z* or *Y* . To this end, we need to adapt the BCR and SFDR to the bivariate case. BCR is motivated to incorporate the information about sign of the parameter, in this case, four scenarios regarding the signs for 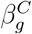 and 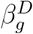 are possible,

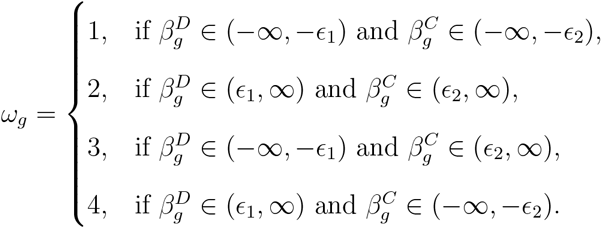

so the bivariate BCR is defined as 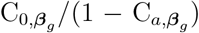 where 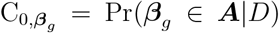 and 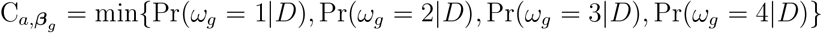. Similarly, mSFDR can be adjusted based on these four cases (expressions are omitted here for conciseness).

We applied BISHOT to the scRNAseq data while incorporating LFCs from bulk RNAseq (denoted as *µ*_*g*_) in the prior, specifically, ***h***_*g*_ = (*κµ*_*g*_, *µ*_*g*_)^⊤^ where *κ* is a parameter potentially used to specify the sign and size of 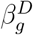 through *µ*_*g*_. With *ϵ*_1_ = *ϵ*_2_ = log_2_(1.5), we identified 436 DE genes at *SFDR* = 0.05. When *ϵ*_1_ = *ϵ*_2_ = log_2_(1.2), this number increased to 2642. For comparison, we applied MAST (Finak et al., 2015) to the data using the `FindMarkers` function in the `R Seurat` package (Hao et al., 2024), which yielded 108 DE genes. Figure 8 left panel presents the overlaps in DE genes across these methods: BISHOT with larger ***ϵ*** = (*ϵ*_1_, *ϵ*_2_)^⊤^, identified 65 out of the 108 DE genes from MAST. For lower ***ϵ***, BISHOT was able to identify almost all the DE genes from MAST. Notably, BISHOT identified 371 additional DE genes, within the intersection of genes identified using both thresholds, that were not identified by MAST.

**Figure 8:**
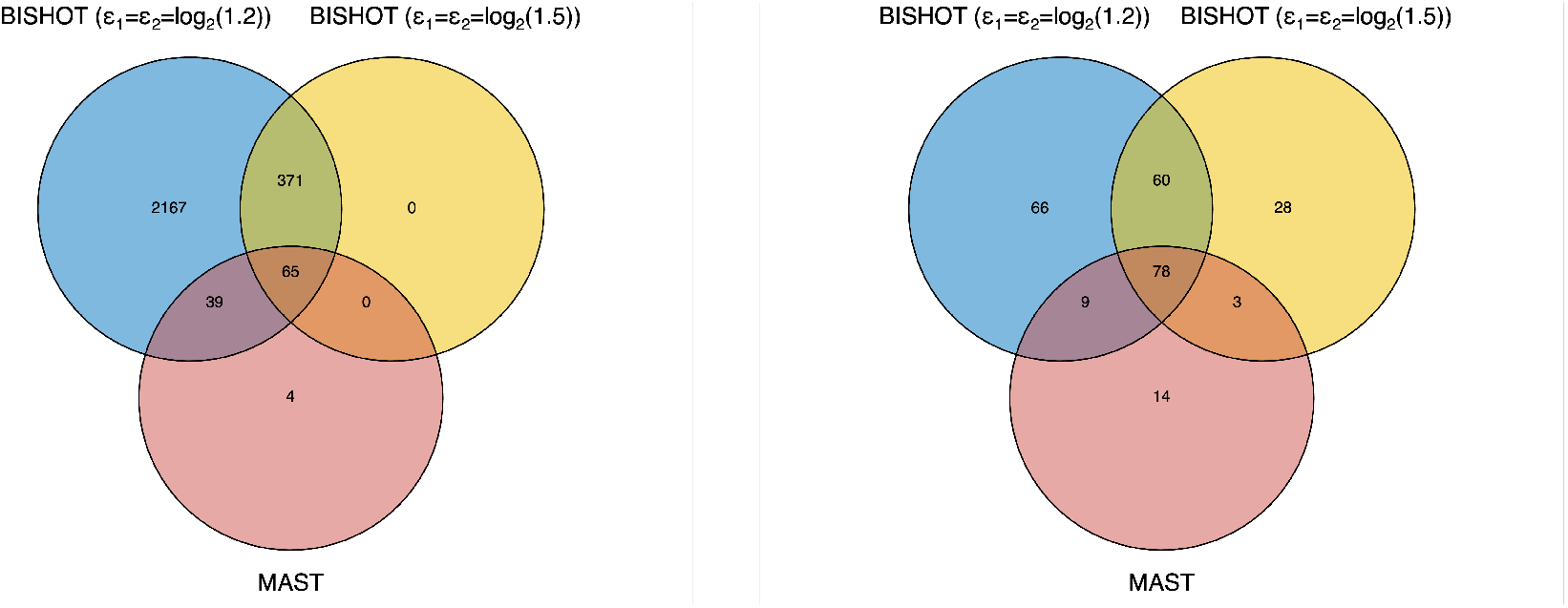
Venn diagrams comparing different methods in terms of identified DE genes (left panel) and enriched GO terms (right panel).

To further investigate the biological interpretations of the DE genes, we performed Gene Ontology (GO) enrichment analysis based on the `GSEA` software (Subramanian et al., 2005). As shown in Figure 8 right panel, there were 60 significant (FDR*<*0.05) GO terms that were identified by BISHOT with both thresholds, but not by MAST. The Appendix B provides a list of these GO terms, many of which are related to lung aging. The endothelial barrier (GO:0061028) is crucial for lung function, regulating substance exchange between blood and tissue. Senescence of endothelial cells during aging compromises this barrier, increasing susceptibility to acute lung injury and acute respiratory distress syndrome (Barabutis et al., 2016; Najari Beidokhti et al., 2025). Mitochondria, essential for energy, senescence, apoptosis, and regeneration of type-2 pneumocytes, show impaired function with age due to altered expression of mitochondrial genes and ribosomal subunits (GO:0140053, GO:0005763, GO:0005762) (Cloonan et al., 2020). Cell morphogenesis (GO:0000902) disruptions contribute to lung aging, marked by alveolar enlargement and structural changes known as the “senile lung” (Wang et al., 2024). Protein serine/threonine kinases (GO:0004674) influence age-related lung disease. For example, PINK1 regulates mitophagy, with its dysregulation linked to fibrosis and COPD (Mizumura et al., 2014). Overexpression of MAP kinase-interacting serine/threonine kinase 2 promotes lung cancer progression (Guo et al., 2017), while protein kinase B (Akt) signaling protects myofibroblast from apoptosis, making it a therapeutic target for lung fibrosis (Wang et al., 2022).

## 7 Discussions

As far as we know there is an absence in the literature where a multiple testing methodology was applied to ***β***_g_ ∈ **R**^*p*^ when *p >* 1. We have seen a success for *p* = 2 in the single cell data application in Section 6 on non-normally distributed data. This conveys a positive signal of the flexibility of BISHOT regarding model and parameter choices and therefore is of interest on a separate note to the multiple testing regime.

We consider a composite null hypothesis in this paper as desired by the practical problems we described. It is observed that the role of *ϵ* is connected with conservativeness in terms of the actual SFDR. More thorough investigation regarding the combined effect of *ϵ*, the true distribution of *β*_*g*_ and the accuracy of *h*_*g*_ (these factors are what we believe contribute to the degree of conservativeness of our method) is beyond the scope of this paper and is left as potentially a future work.

BISHOT offers a general framework for Bayesian differential analysis that incorporates prior knowledge with relatively low computational cost and remarkable accuracy. In this paper, we illustrate the framework using normal linear and two-component models, though it is broadly applicable to other models. Extending BISHOT to other frequently used models, such as the negative binomial model (Robinson et al., 2010; Wu et al., 2013), is a direction for future research.

In this work, we limit ourselves to the specifications of the shrinkage parameters *λ*_*g*_, *τ* based on the original horseshoe. As we explained in Section 5 there exists a close proximity between the notion of sparsity (in nonzero parameters) and the consistency between *h*_*g*_ and 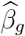. Given the well-known drawbacks for the horseshoe prior distributions, we can extend the proposed framework to be combined with more recent developments regarding the horseshoe prior (Bhadra et al., 2017), in order to tackle the various scenarios regarding the overall consistency between the two studies.

## Acknowledgements

High Performance Computing resources provided by the High Performance Research Computing (HPRC) core facility at Virginia Commonwealth University (https://hprc.vcu.edu) were used for conducting the research reported in this work. This research was supported by the Biostatistics and Bioinformatics Shared Resource of the University of Kentucky Markey Cancer Center (P30CA177558).

## Data availability

The AML data (Hohmann et al., 2016) used in Section 5 is available from GEO under accession number GSE79284. The lung aging data (Angelidis et al., 2019) used in Section 6 is available from Zenodo at https://doi.org/10.5281/zenodo.5048449, which is provided by Squair et al. (2021).

## Supporting Information

Appendix A referenced in Sections 2 and 4 and Appendix B, referenced in Sections 5 and 6, is available with this paper at the journal website.

