## Supplementary Material for "A Bayesian Informative Shrinkage Approach for Large-scale Multiple Hypothesis Testing (BISHOT): with Applications in Differential Analysis of Omics Data"

#### S.1 Appendix A

##### S.1.1 Proof of Theorem 1

We first calculate the expected value of $L_{1,\lambda}(\boldsymbol{\delta}, \boldsymbol{\omega})$ under the posterior distribution of $p(\boldsymbol{\omega}|D)$:

$$E_{\boldsymbol{\omega}|D}\{L_{1,\lambda}(\boldsymbol{\delta}, \boldsymbol{\omega})\} = \frac{1}{G}\sum_{g=1}^{G}\{\lambda\widetilde{\omega}_{0g}\delta_g + \widetilde{\omega}_{1g}(1-\delta_g)\}.$$

The optimal choices for $\delta_g$ would be $\delta_g = I\{\lambda\widetilde{\omega}_{0g} < \widetilde{\omega}_{1g}\}$.

Repeat the above process for $L_{2,\lambda}$, the optimal choices for $\delta_g$ corresponding to $L_{2,\lambda}$ is $\delta_g = I\{\lambda\widetilde{\omega}_{0g} < \widetilde{\omega}_{2g}\}$.

##### S.1.2 Proof of Theorem 2

The proof is similar to that of Theorem 1.

##### S.1.3 Proof of Theorem 3

The proof extends that of Theorem 1 in Fu et al. (2022) in the framework of the proposed Bayesian procedure. We first prove two properties about $\alpha(\lambda) = \mathrm{SFDR}(\boldsymbol{\delta}^{OPT})$: (i) $\alpha(\lambda) < \lambda$ and (ii) $\alpha(\lambda)$ is non-decreasing.

For any $\alpha$ given, suppose $\lambda$ is chosen so that $\mathrm{mSFDR}(\boldsymbol{\delta}^{OPT}) = \alpha$. For an arbitrary decision $\boldsymbol{d} = (d_1, \ldots, d_G)$ satisfying $\mathrm{SFDR}(\boldsymbol{d}) \leq \alpha$. Equivalently,

$$
\begin{aligned}
E\bigg(\sum_{g\in\mathcal{S}}\{(1-\alpha)\widetilde{\omega}_{0g} - \alpha\widetilde{\omega}_{1g}\}\delta_g^{OPT} + \sum_{g\in\mathcal{S}^c}\{(1-\alpha)\widetilde{\omega}_{0g} - \alpha\widetilde{\omega}_{2g}\}\delta_g^{OPT}\bigg) &= 0 \\
E\bigg(\sum_{g\in\mathcal{S}}\{(1-\alpha)\widetilde{\omega}_{0g} - \alpha\widetilde{\omega}_{1g}\}d_g + \sum_{g\in\mathcal{S}^c}\{(1-\alpha)\widetilde{\omega}_{0g} - \alpha\widetilde{\omega}_{2g}\}d_g\bigg) &\leq 0.
\end{aligned}
$$

Immediately,

$$
E\bigg(\sum_{g\in\mathcal{S}}\{(1-\alpha)\widetilde{\omega}_{0g} - \alpha\widetilde{\omega}_{1g}\}(\delta_g^{OPT} - d_g) + \sum_{g\in\mathcal{S}^c}\{(1-\alpha)\widetilde{\omega}_{0g} - \alpha\widetilde{\omega}_{2g}\}(\delta_g^{OPT} - d_g)\bigg) \geq 0. \quad \text{(S.1)}
$$

Let $t = (\lambda - \alpha)/(1-\lambda) > 0$ (from (i)). If $g \in \mathcal{S}$, when $\delta_g^{OPT} = 1$, we have $\widetilde{\omega}_{0g} < \lambda(\widetilde{\omega}_{0g} + \widetilde{\omega}_{1g})$, equivalently, $t\widetilde{\omega}_{1g} > (1-\alpha)\widetilde{\omega}_{0g} - \alpha\widetilde{\omega}_{1g}$, otherwise when $\delta_g^{OPT} = 0$, $t\widetilde{\omega}_{1g} \leq (1-\alpha)\widetilde{\omega}_{0g} - \alpha\widetilde{\omega}_{1g}$. Hence $\sum_{g\in\mathcal{S}}\{(1-\alpha)\widetilde{\omega}_{0g} - \alpha\widetilde{\omega}_{1g}\}(\delta_g^{OPT} - d_g) \leq t\sum_{g\in\mathcal{S}}\widetilde{\omega}_{1g}(\delta_g^{OPT} - d_g)$. Similarly, $\sum_{g\in\mathcal{S}^c}\{(1-\alpha)\widetilde{\omega}_{0g} - \alpha\widetilde{\omega}_{2g}\}(\delta_g^{OPT} - d_g) \leq t\sum_{g\in\mathcal{S}^c}\widetilde{\omega}_{2g}(\delta_g^{OPT} - d_g)$. Together with (S.1),

$$
E\bigg(\sum_{g\in\mathcal{S}}\widetilde{\omega}_{1g}(\delta_g^{OPT} - d_g) + \sum_{g\in\mathcal{S}^c}\widetilde{\omega}_{2g}(\delta_g^{OPT} - d_g)\bigg) \geq 0.
$$

This indicates that $\mathrm{SETP}(\boldsymbol{\delta}^{OPT}) \geq \mathrm{SETP}(\boldsymbol{d})$ and concludes the proof.

##### S.1.4 Connection of BCR to Bayes factor

Recall $\Theta_0 = (-\epsilon, \epsilon)$, $\Theta_1 = (-\infty, -\epsilon)$ and $\Theta_2 = (\epsilon, \infty)$. Let $\Pi|_{\Theta_i}(\cdot) = \Pi(\cdot)/\Pi(\Theta_i)$ be the prior distribution $\Pi$ restricted on $\Theta_i$ $(i = 0, 1, 2)$, and the marginal distribution of data under $H_i$ is $\Pr(D|H_i) = \int_{\beta\in\Theta_i}\Pi|_{\Theta_i}(\beta)P(D|\beta)d\beta$ $(i = 0, 1, 2)$, therefore

$$
\mathrm{BF}_{i0} = \frac{\Pr(D|H_i)}{\Pr(D|H_0)} = \frac{\int_{\beta\in\Theta_i}\Pi(\beta)\Pr(D|\beta)d\beta}{\int_{\beta\in\Theta_0}\Pi(\beta)\Pr(D|\beta)d\beta}\frac{\Pi(\Theta_0)}{\Pi(\Theta_i)} \text{ for } i = 1, 2.
$$

Without loss of generality, let us assume that $\Pi(\Theta_1) = \Pi(\Theta_2)$. From the expression of BCR in Section 2.2, we immediately find that $C_\beta = 1/(1 + \Pi(\Theta_1) \cdot \Pi(\Theta_0)^{-1} \cdot \max\{\mathrm{BF}_{10}, \mathrm{BF}_{20}\})$.

##### S.1.5 Conditional distributions of parameters in models (1) and (3)

$$
\begin{aligned}
&\Pr(c_g, \beta_g | \lambda_g, \tau, D_g) \sim \mathrm{t}_{\nu_g}(\mu_g, s_g) \\
&\Pr(\lambda_g \, | \beta_g, \tau, \sigma_g^2, h_g) \propto \frac{1}{\lambda_g(1+\lambda_g^2)} \exp\left\{ - \frac{(\beta_g - h_g)^2}{2\lambda_g^2 \tau^2 \sigma_g^2} \right\} I(\lambda_g > 0) \\
&\Pr(\tau \, | \beta_1, \ldots, \beta_G, \sigma_1^2, \ldots, \sigma_G^2, h_1, \ldots, h_G) \propto \frac{1}{(1+\tau^2)\tau^G} \exp\left\{ - \sum_{g=1}^{G} \frac{(\beta_g - h_g)^2}{2\lambda_g^2 \tau^2 \sigma_g^2} \right\} I(\tau > 0) \\
&\Pr(\sigma_g^2 | \beta_g, \lambda_g, \tau, h_g) \sim \mathrm{IG}(\widetilde{\xi}_1, \widetilde{\xi}_2),
\end{aligned}
$$

where $\mu_g = Q_g^{-1}\eta_g$ with $Q_g = \sum_i \widetilde{X}_i \widetilde{X}_i^\top + \mathrm{diag}(100^{-1}, \lambda_g^{-2}\tau^{-2})$ and $\eta_g = \sum_i \widetilde{X}_i Y_{ig} + \lambda_g^{-2}\tau^{-2}h_g$, $s_g = Q_g^{-1}\{\xi_1 + n/2\}^{-1}\{\xi_2 + (\sum_i Y_{ig}^2 + \lambda_g^{-2}\tau^{-2}h_g^2 - \eta_g^\top Q_g^{-1}\eta_g)/2\}$ and $\nu_g = 2\xi_1 + n$. $\mathrm{t}_\nu(\mu, s)$ denotes a multivariate t distribution with center $\mu$ and scale $s$ and degrees of freedom $\nu$, $\widetilde{\xi}_1 = \xi_1 + n/2 + 1$, $\widetilde{\xi}_2 = \xi_2 + (c_g^2/10^2 + (\beta_g - h_g)^2\tau^{-2}\lambda_g^{-2} + \sum_i (Y_{ig} - c_g - X_i^\top \beta_g)^2)/2$. Here $\widetilde{X}_i$ is $(1, X_i^\top)^\top$ which expands $X_i$ with a constant 1.

##### S.1.6 Simulation II and results

In simulation II, we adopt the same mechanism for the set of DE genes $\Omega$ but try to explore a scenario when the DE gene differential expressions has a unimodal distribution at zero, which indicates that true DE expressions could be close to zero rather than a distance away from zero. Specifically, if $g \in \Omega$, $\beta_g \sim 0.4\mathrm{N}(0, 0.25^2) + 0.2\mathrm{N}(0, 0.5^2) + 0.2\mathrm{N}(0, 1^2) + 0.2\mathrm{N}(0, 2^2)$. This DE distribution is the same as the spiky scenario in Stephens (2017).

The parameters needed in model (1) are set as follows: $c_g = 4$, $\sigma_g = 1.5$ for a total $G = 5000$ genes. $\epsilon$ is set to be $\epsilon = 0.01$ or $\epsilon = 0.05$ and $h_g \sim \mathrm{N}(\beta_g, 0.1^2)$ in reflection of the size of the true DE genes generated above ($\epsilon$ and $h_g$ should be set to be adaptive to the distribution of true DE genes practically). The remaining parameter/hyperparameter choices are identical to the description in Section 4. See Figures S.1-S.4 for the similar analyses comparing BISHOT and SunSpike0 as presented in Section 4 based on 50 simulated data sets.

As observed in Section 4, BISHOT is more robust among all scenarios spanned by $p$ and $n$ in producing SFDR estimates and accuracy in top ranked genes. BISHOT appears to produce some "fake" discoveries when $\epsilon$ is larger and the SFDR nominal threshold is small (right panel in Figure S.2). After close examination these are actual nonzero $\beta_g$ that are smaller than but close to $\epsilon$, whose identification are caused by their priors sticking out. This shall suggest that we select $\epsilon$ appropriately by not neglecting the bulk of the distribution of $\beta_g$, e.g. the middle probability corresponding to $|\beta_g| < \epsilon (= 0.05)$ is about 10% which, when combined with a relative large prior $h_g$, can possibly lead to the aforementioned "fake" discoveries.

### S.2 Appendix B

#### S.2.1 Venn diagram comparing DE results from BISHOT and SunSpike0 in Section 5

#### S.2.2 Enriched GO terms from the DE analysis in Section 6

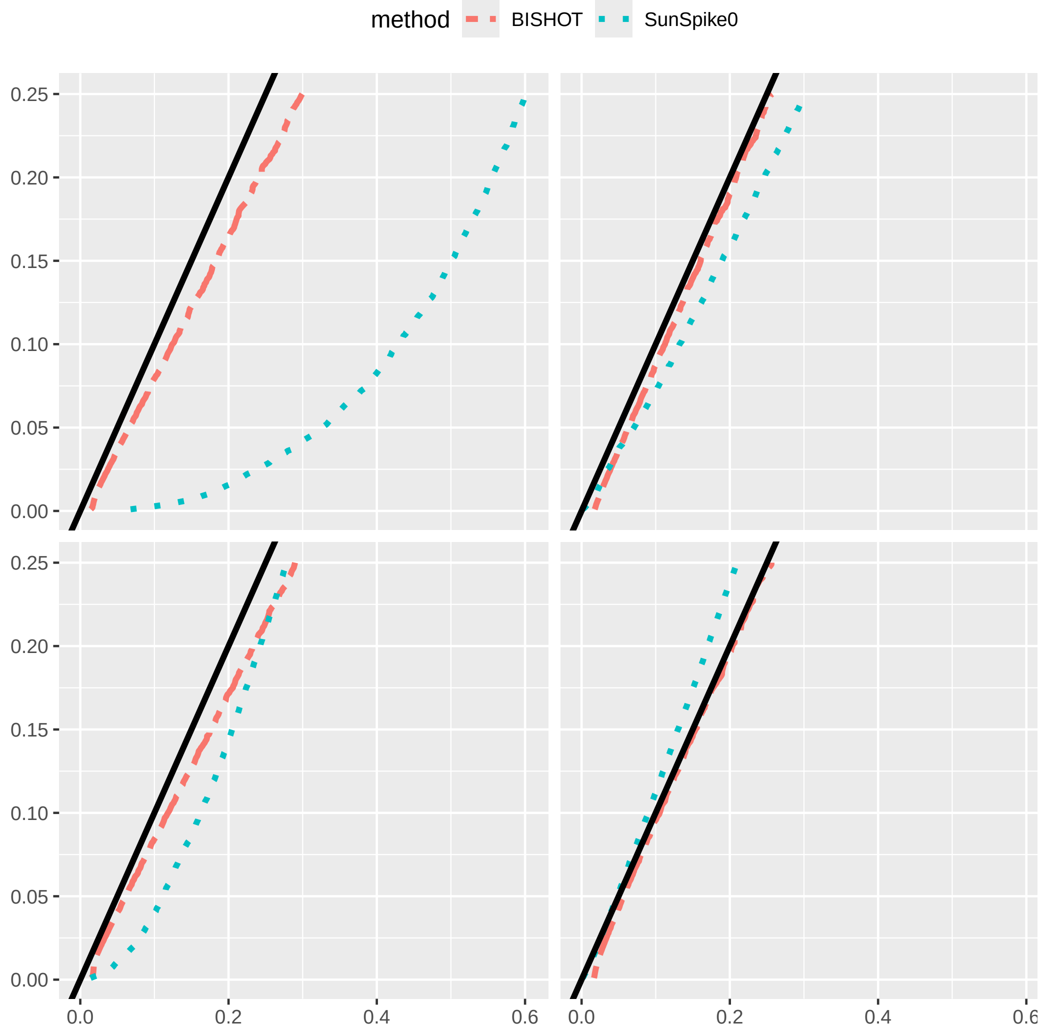


Figure S.1: The actual signed false discovery rate (SFDR) when $\epsilon = 0.01$ averaged over 50 simulated data using mSFDR threshold 0.001 to 0.25 corresponding to 10 (left panel) or 50 (right panel) samples in the treatment and control group. The top and bottom panel displays the case when $p = 0.1$ and $p = 0.5$.

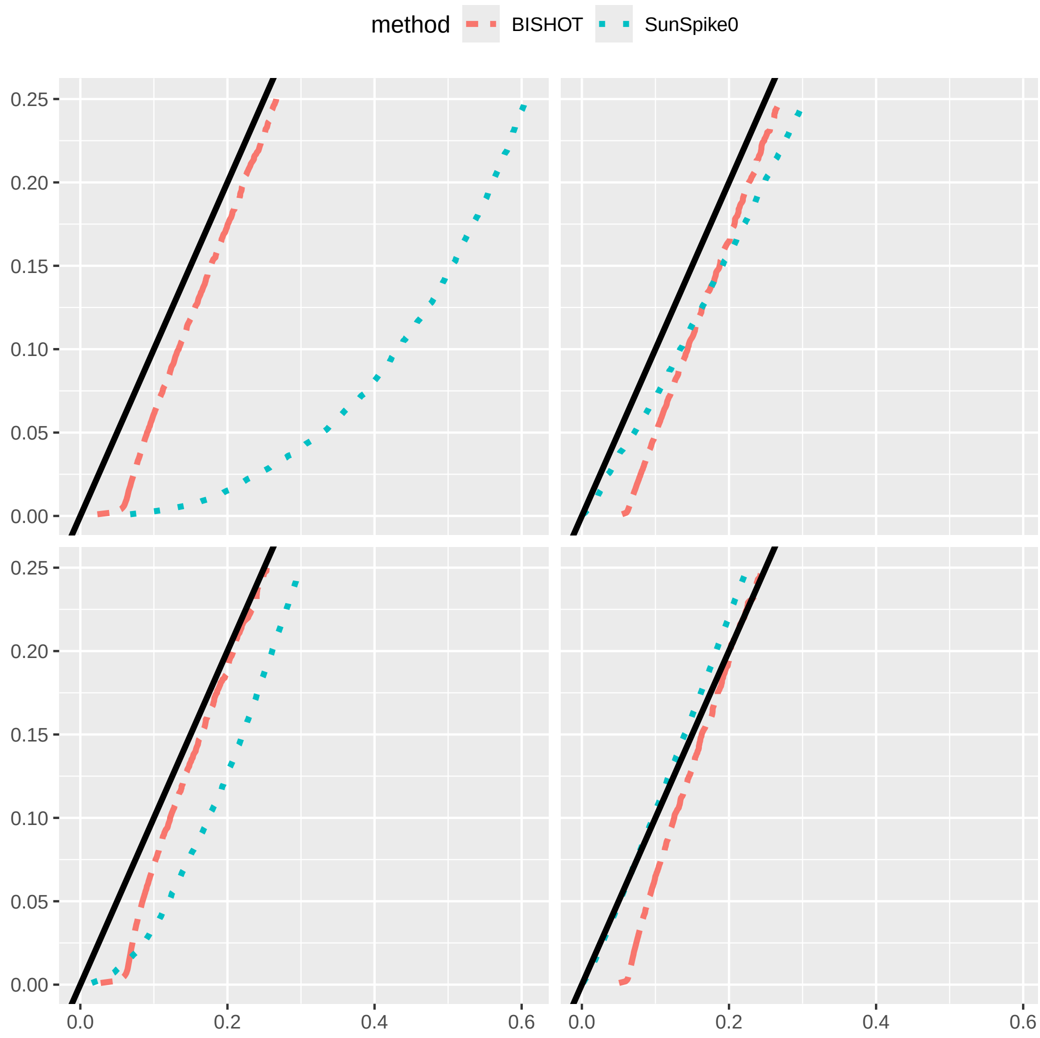


Figure S.2: The actual signed false discovery rate (SFDR) when $\epsilon = 0.05$ averaged over 50 simulated data using mSFDR threshold 0.001 to 0.25 corresponding to 10 (left panel) or 50 (right panel) samples in the treatment and control group. The top and bottom panel displays the case when $p = 0.1$ and $p = 0.5$.

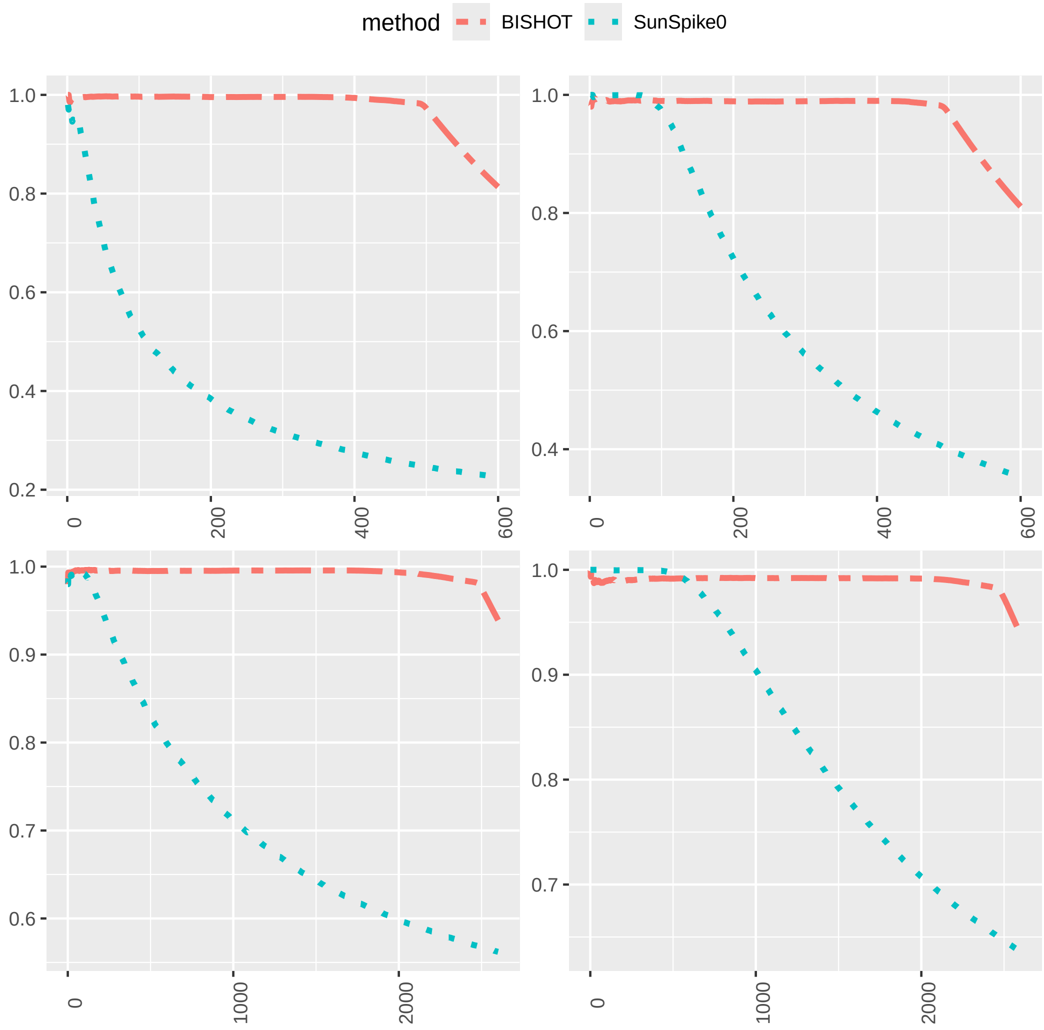


Figure S.3: The proportion of true DE genes for the first $R$ genes with smallest $q$ values when $\epsilon = 0.01$ averaged over 50 simulated data for $R = 1, \ldots, 5000p+100$ corresponding to 10 (left panel) or 50 (right panel) samples in the treatment and control group. The top and bottom panel displays the case when $p = 0.1$ and $p = 0.5$.

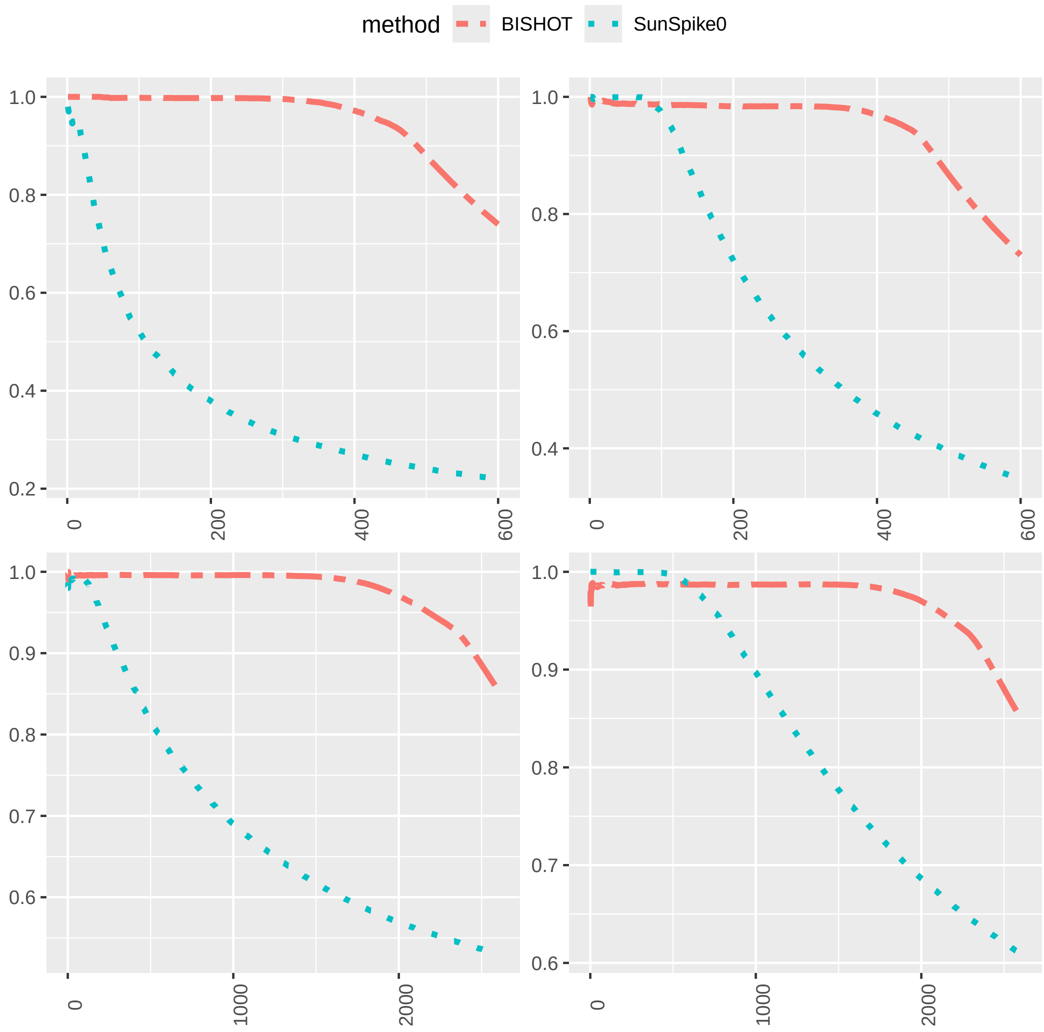


Figure S.4: The proportion of true DE genes for the first $R$ genes with smallest $q$ values when $\epsilon = 0.05$ averaged over 50 simulated data for $R = 1, \ldots, 5000p+100$ corresponding to 10 (left panel) or 50 (right panel) samples in the treatment and control group. The top and bottom panel displays the case when $p = 0.1$ and $p = 0.5$.

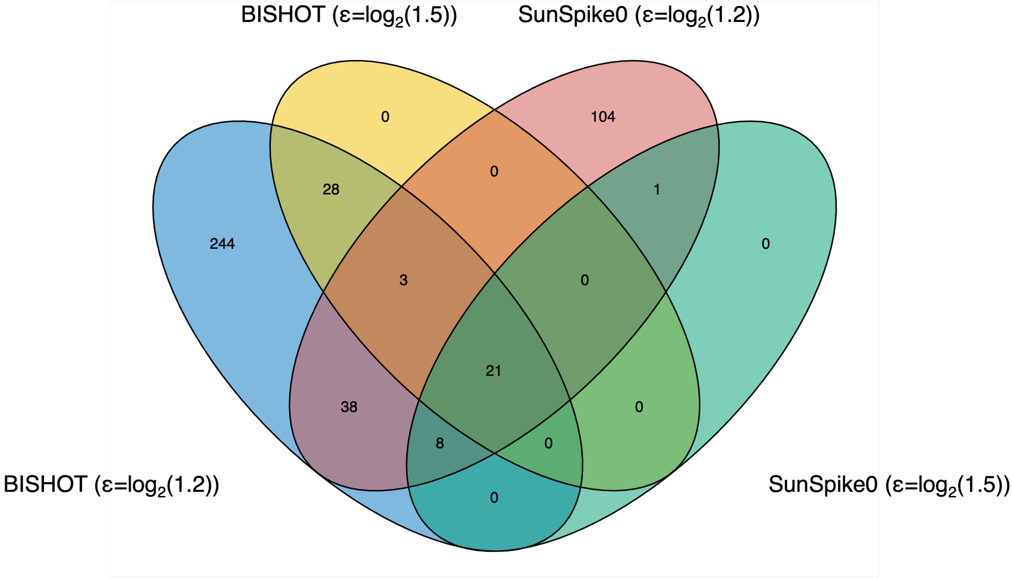


Figure S.5: Venn diagram comparing DE genes identified by different methods from the DE analysis in Section 5.

| GO Term ID | Term Name | Category | Alteration |
|---|---|---|---|
| GO:0000902 | cell morphogenesis | Biological Process | up-regulated |
| GO:0004674 | protein serine/threonine kinase activity | Molecular Function | up-regulated |
| GO:0030010 | establishment of cell polarity | Biological Process | up-regulated |
| GO:0004672 | protein kinase activity | Molecular Function | up-regulated |
| GO:0002399 | spleen development | Biological Process | up-regulated |
| GO:0045669 | positive regulation of cell differentiation | Biological Process | up-regulated |
| GO:0048589 | developmental growth | Biological Process | up-regulated |
| GO:0004674 | protein serine/threonine kinase activity | Molecular Function | up-regulated |
| GO:0008361 | regulation of cell size | Biological Process | up-regulated |
| GO:0048638 | regulation of developmental growth | Biological Process | up-regulated |
| GO:0061028 | establishment of endothelial barrier | Biological Process | up-regulated |
| GO:0010494 | cytoplasmic stress granule | Cellular Component | up-regulated |
| GO:0007017 | microtubule-based process | Biological Process | up-regulated |
| GO:0045071 | negative regulation of viral genome replication | Biological Process | up-regulated |
| GO:0003730 | mRNA 3'-UTR binding | Molecular Function | up-regulated |
| GO:0060560 | developmental growth involved in morphogenesis | Biological Process | up-regulated |
| GO:0009790 | embryo development | Biological Process | up-regulated |
| GO:0003682 | chromatin binding | Molecular Function | up-regulated |
| GO:0009887 | animal organ morphogenesis | Biological Process | up-regulated |
| GO:0004812 | aminoacyl-tRNA ligase activity (aminoacyltransferase activity) | Molecular Function | up-regulated |
| GO:0048471 | perinuclear region of cytoplasm | Cellular Component | up-regulated |
| GO:0048588 | developmental cell growth | Biological Process | up-regulated |
| GO:0008610 | lipid biosynthetic process | Biological Process | up-regulated |
| GO:0001223 | transcription coactivator binding | Molecular Function | up-regulated |
| GO:0097542 | ciliary tip | Cellular Component | up-regulated |
| GO:0040007 | growth | Biological Process | up-regulated |
| GO:0044782 | cilium organization | Biological Process | up-regulated |
| GO:0060041 | retina morphogenesis in camera-type eye | Biological Process | up-regulated |
| GO:0009792 | embryo development ending in birth or egg hatching | Biological Process | up-regulated |
| GO:0001570 | vasculogenesis | Biological Process | up-regulated |
| GO:0051020 | GTPase binding | Molecular Function | up-regulated |
| GO:0006412 | translation | Biological Process | down-regulated |
| GO:0005762 | mitochondrial large ribosomal subunit | Cellular Component | down-regulated |
| GO:0022613 | ribonucleoprotein complex biogenesis | Biological Process | down-regulated |
| GO:0140053 | mitochondrial gene expression | Biological Process | down-regulated |
| GO:0002181 | cytoplasmic translation | Biological Process | down-regulated |
| GO:0016469 | proton-transporting two-sector ATPase complex | Cellular Component | down-regulated |
| GO:0042254 | ribosome biogenesis | Biological Process | down-regulated |
| GO:0050819 | regulation of coagulation | Biological Process | down-regulated |
| GO:0005791 | rough endoplasmic reticulum | Cellular Component | down-regulated |
| GO:0005761 | mitochondrial small ribosomal subunit | Cellular Component | down-regulated |
| GO:1990908 | endonuclease complex | Cellular Component | down-regulated |
| GO:0061077 | chaperone-mediated protein folding | Biological Process | down-regulated |
| GO:0009141 | nucleoside triphosphate metabolic process | Biological Process | down-regulated |
| GO:0042274 | ribosomal small subunit biogenesis | Biological Process | down-regulated |
| GO:0006457 | protein folding | Biological Process | down-regulated |
| GO:0000955 | RNA 5'-end processing | Biological Process | down-regulated |
| GO:0005771 | multivesicular body | Cellular Component | down-regulated |
| GO:0006364 | rRNA processing | Biological Process | down-regulated |
| GO:0032040 | small-subunit processome | Cellular Component | down-regulated |
| GO:0031970 | organelle envelope lumen | Cellular Component | down-regulated |
| GO:0051082 | unfolded protein binding | Molecular Function | down-regulated |
| GO:0016072 | rRNA metabolic process | Biological Process | down-regulated |
| GO:0044183 | protein folding chaperone | Molecular Function | down-regulated |
| GO:0007006 | mitochondrial membrane organization | Biological Process | down-regulated |
| GO:0051087 | protein folding chaperone binding | Molecular Function | down-regulated |
| GO:0032509 | endosome transport via multivesicular body sorting pathway | Biological Process | down-regulated |
| GO:0097525 | spliceosomal tri-snRNP complex | Cellular Component | down-regulated |
| GO:0071985 | multivesicular body sorting pathway | Biological Process | down-regulated |
| GO:0006397 | mRNA processing | Biological Process | down-regulated |

Figure S.6: A list of enriched GO terms identified by BISHOT from the DE analysis in Section 6.
